# Wnt-presenting materials sustain H3K14-acetylation in human skeletal stem cells for tissue engineering and bone repair

**DOI:** 10.64898/2025.12.18.694408

**Authors:** Pierre Becquart, Pierre Tournier, Sergi Junyent, Binbin Ma, Yu Liu, Yin Tang, Xin Chen, Dong Fang, Anjali P Kusumbe, Shukry J Habib

**Affiliations:** Department of Biomedical Sciences, University of Lausanne, Rue du Bugnon 7a, CH-1005 Lausanne, Switzerland; California Institute of Technology, Pasadena, CA 91125, USA; Howard Hughes Medical Institute, Department of Biology, Johns Hopkins University, Baltimore, MD, USA; Zhejiang Provincial Key Laboratory for Cancer Molecular Cell Biology, Life Sciences Institute Zhejiang University, Zhejiang, China; Department of Ministry of Education, The Second Affiliated Hospital, Zhejiang University School of Medicine, Zhejiang, China; Department of Biology, Johns Hopkins University, Baltimore, MD, USA; Tissue and Tumor Microenvironments Laboratory, Cancer Discovery and Regenerative Medicine Program, Lee Kong Chian School of Medicine, Nanyang Technological University, Singapore

## Abstract

Engineering functional tissues for transplantation requires insight into epigenetic mechanisms that regulate stem cell fate. We have developed the Wnt-induced osteogenic tissue model (WIOTM), a platform that recapitulates human osteogenesis, and identified acetylation of histone H3 at lysine 14 (H3K14ac) as a critical epigenetic regulator in human skeletal stem cells (hSSCs). In WIOTM, localized Wnt signals drive asymmetric cell division (ACD), yielding a proximal hSSC with high H3K14ac and a distal daughter with reduced H3K14ac that migrates into the 3D collagen matrix and initiates osteogenic differentiation. Disrupting H3K14ac in hSSCs abrogates ACD and WIOTM formation. To test whether hSSCs maintain H3K14ac *in vivo*, we formed the WIOTM on Wnt-functionalized polymer bandages and transplanted them into calvarial defects. The WIOTM contributed to bone repair, and human cells adjacent to the bandages retained high H3K14ac despite the injury environment. These findings establish WIOTM as both a mechanistic and translational platform for regenerative medicine.

## Introduction

Tissue engineering has transformed the field of regenerative medicine, instilling hope for treating previously incurable injuries and diseases. By replicating the fundamental principles of tissue formation, tissue engineering creates functional substitutes for damaged or lost tissues^1^. Stem cells play a critical role in tissue engineering, as they possess the unique ability to differentiate into the required cell types and thereby promote tissue formation^2^.

Asymmetric cell division is a mechanism that regulates mammalian stem cells by producing one self-renewed stem cell, and another daughter cell which contributes to the pool of differentiated cells during tissue formation. In many tissues, including skeletal tissues, maintaining stem and progenitor cell pools is essential not only for effective regeneration but also for sustaining tissue health with aging^3,4^. Wnt proteins are key mediators of asymmetric cell division^1,2,5–7^. Wnt ligands are hydrophobic and often act via localized secretion and presentation. Activated Wnt receptors in the responsive cell promote β-catenin translocation to the nucleus and the transcription of genes that regulate many aspects of cell function and fate^2,4^.

We developed synthetic surfaces with covalently bound Wnt ligands that replicate the effects of localized Wnt presentation^6,8,9^. Using these surfaces, we established the Wnt-induced osteogenic tissue model (WIOTM), a 3D collagen-based organoid in which hSSCs are seeded on immobilized Wnt proteins and overlaid with a type I collagen gel and osteogenic differentiation medium^9^ (Figure 1A). During cell division, Wnt-responsive hSSCs orient their spindle perpendicularly to the Wnt surface, asymmetrically segregating stem cell and osteogenic markers^10^. One daughter cell remains in contact with the Wnt signal and self-renews, while the Wnt-distal cell commits to osteogenic differentiation and migrates into the collagen gel. This generates a stable hSSC population adjacent to the Wnt source and a gradient of increasingly differentiated osteogenic progeny through the gel’s thickness. The WIOTM recapitulates key aspects of human osteogenesis and the skeletal stem cell niche, providing a powerful model for dissecting mechanisms of asymmetric cell division in human cells.

**Figure 1:**
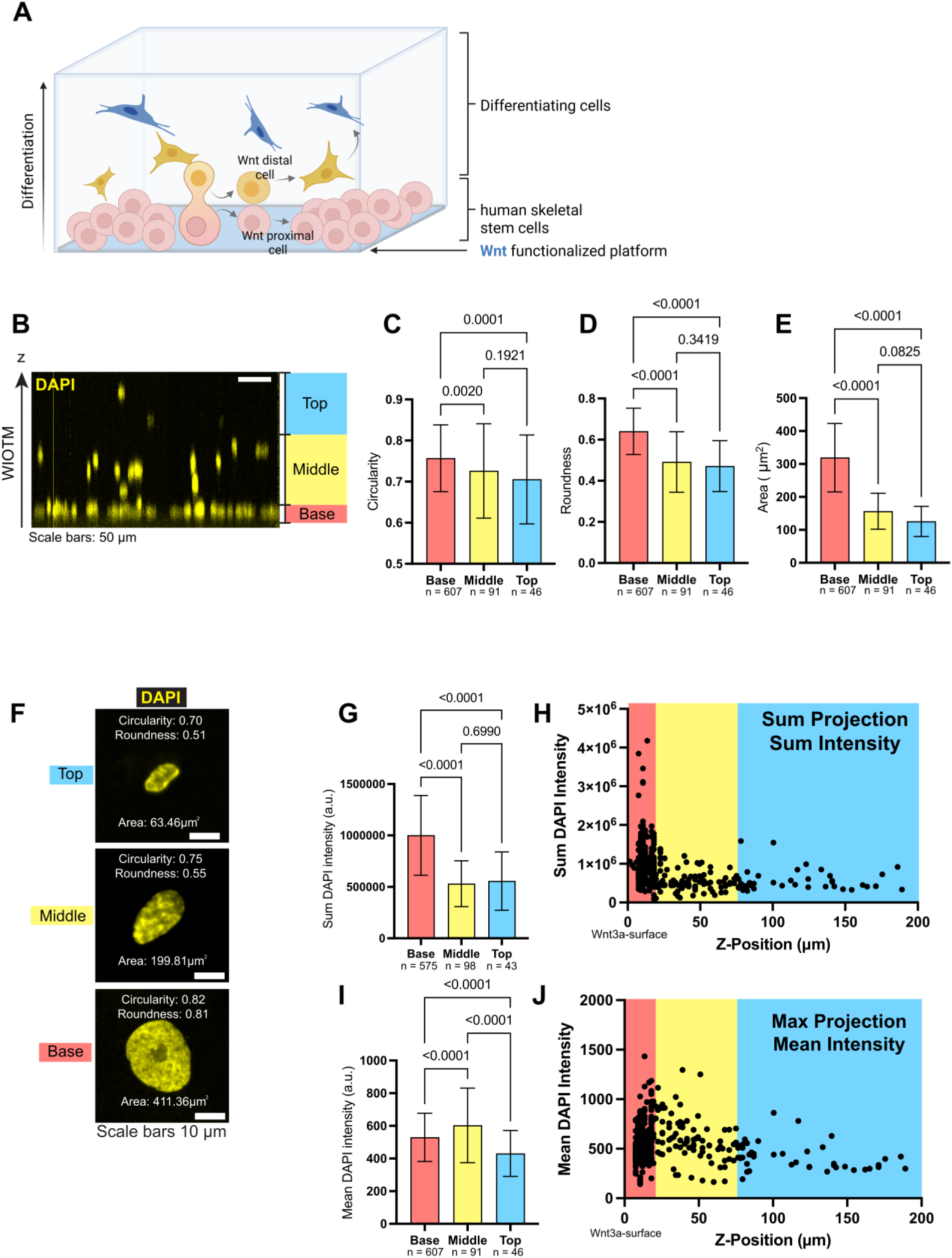
Nuclear morphology of human skeletal stem cells (hSSCs) changes as cells migrate in the WIOTM. (A) Schematic representation of the WIOTM. hSSCs are seeded on a Wnt functionalized platform, covered with a gel of collagen I and osteogenic differentiation media. After 48 h, the first asymmetric cell division can be observed, giving rise to two daughter cells, one retaining its stem cell identity (Wnt proximal cell; red cell), the other one starting differentiation processes (Wnt distal cell; yellow cell). After 7 days of culture, differentiating cells are migrating into the gel and further differentiate towards the osteogenic lineage (blue cells). (B) Representative side-view (z axis) projection image of hSSC migration within the WIOTM after 7 days in culture. Nuclei are stained with DAPI (yellow). The WIOTM is segmented into three layers to assess cell migration depth: Base (red; 0–10 μm), Middle (yellow; 10–75 μm), and Top (blue; above 75 μm). (C–E) Quantification of nuclear morphology based on DAPI staining across the three WIOTM layers: (C) circularity, (D) roundness, and (E) area. (F) Representative images of DAPI-stained nuclei in each WIOTM layer, illustrating morphological differences. (G–J) Quantitative analysis of DAPI signal intensity per cell in each WIOTM layer. (G–H) Measurements derived from summed projection images and total sum DAPI intensity per cell. (I–J) Measurements from maximum projection images and mean DAPI intensity per cell. Data in panels (C, D, E, G, I) are presented as mean ± standard deviation (s.d.). Statistical comparisons were performed using uncorrected Fisher’s LSD test. Sample sizes n (cell) are indicated within each graph.

Initial asymmetric distribution of proteins in daughter cells must be further sustained to ensure diverging cell fates via differing transcriptional programs. Epigenetic regulation of gene expression is influenced not only by the formation of transcriptional complexes (e.g. β-catenin-TCF/LEF), but also by genomic DNA packaging. For example, lysine residues of Histone 3 (H3K) can exist in various states of post-translational acetylation which alters local packaging of chromatin, and so controls the accessibility to transcriptional machinery and resulting gene expression^11,12^. In hSSCs, histone acetylation is implicated in differentiation^13–15^, including H3 acetylation in osteogenesis^16–20^, which can be influenced through the histone acetyl transferases and/or the inhibition of histone deacetylases. While these studies provide valuable insights into molecular mechanisms, they are often conducted in 2D cell culture models that inadequately represent the spatial, mechanical, and cellular heterogeneity of 3D tissues. Therefore, little is known about H3 lysine acetylation during mammalian asymmetric stem cell division and human osteogenic tissue formation. Particularly, the role that biomaterials can play in influencing epigenetic modifications of hSSCs in 3D cultures and *in vivo* is poorly understood. We previously developed WIOTM-bandages using polycaprolactone-based (PCL) material, which we showed can expedite the regeneration of mature, well-organized bone after inducing a critical size defect in mouse models. Notably, hSSCs were maintained close to the bandage and generated osteogenic progenies that contributed to the formation of new bone^10^.

In this study, we identify histone H3 lysine 14 acetylation (H3K14ac) as a key regulator of Wnt-driven asymmetric cell division and WIOTM formation. Genome-wide analyses further show that localized Wnt signaling induces broad chromatin remodeling and activates osteogenic regulatory elements, reinforcing transcriptional programs required for skeletal lineage commitment. We show that immobilized Wnt ligands on various biocompatible polymers support WIOTM formation, which can then be transplanted to promote bone repair. Importantly, transplanted hSSCs adjacent to WIOTM bandages maintained high H3K14ac despite the complexity of the bone injury environment. These findings establish the WIOTM as both a mechanistic model for dissecting asymmetric cell division and tissue formation, and a translational platform for regenerative medicine.

## Results

### Distinct nuclear morphologies across the WIOTM accompany gradients of stemness and osteogenic commitment

The WIOTM, generated by culturing hSSCs on immobilized Wnt proteins overlaid with collagen and osteogenic medium, produces a spatial gradient of stem and osteogenically committed cells that models key features of the skeletal stem cell niche *in vivo*. To investigate how this spatial organization relates to nuclear architecture, we examined nuclear morphology across the WIOTM, reasoning that these differences might reflect underlying changes in chromatin organization and epigenetic state. We initially identified three distinct regions in the WIOTM based on differences in nuclear morphology: Base, where cells contact the Wnt3a; Middle, up to 75 μm; and Top, the region above 75 μm (Figure 1B). Analysis of DAPI staining revealed that hSSCs in contact with the Wnt surface (Base) had larger and less elongated nuclei (Figure 1B-F) with increased total DAPI signal compared to differentiating cells in the Middle and Top regions (Figure 1 G-H). Mean nuclear DAPI intensity varied across regions, with cells in the Middle region displaying higher average intensity than those at the Base or Top (Figure 1 I,J).

Nuclear size is known to be linked to changes in DNA amount, cell cycle stage, chromatin structure, cellular responses to mechanical cues, and transcriptional activity^21^. Furthermore, nuclear shape can be influenced by chromatin packaging^22^. As DNA condensation is crucial for regulation of transcription and influenced by epigenetic marks^23^, we aimed next to investigate variations of histone modifications across the WIOTM. We also wanted to determine whether these variations are established early on and influenced by Wnt-mediated asymmetric stem cell division.

### Localized Wnt signaling drives asymmetric distribution of H3K14ac in hSSC daughter cells and WIOTM regions

Histone H3 (H3) is an evolutionarily conserved DNA-binding protein critical for chromatin packaging and transcriptional regulation ^24^. To examine the role of epigenetic modifications in osteogenesis, we focused on H3 acetylation (ac) marks linked to transcriptional regulation: H3K9ac, H3K14ac, and H3K27ac^16^. H3K9ac and H3K27ac are established markers of active transcription^13,25^, while H3K14ac enrichment at specific promoter regions can prime genes for stimulus-dependent activation in embryonic stem cells^26^. In human skeletal stem cells (hSSCs) and other models, H3K14ac is linked to tissue-specific gene expression^27,28^. We first asked whether chromatin condensation, transcriptional activity, and histone H3 acetylation are established through asymmetric cell division and whether they differ between daughter cells. In pairs of daughter cells derived from hSSC division, we quantified DAPI, RNA polymerase II phosphorylated on Ser2 (RNApolS2P, a marker of active transcription elongation), total H3, H3K9ac, H3K14ac, and H3K27ac. The sum of DAPI intensity was significantly higher in the Wnt3a-contact cell “proximal cell” (Figure 2A-C), consistent with a reduction in total nuclear DAPI staining during osteogenic differentiation in the WIOTM (Figure 1). Total H3, H3K9ac, and H3K27ac were similar between daughter cells. However, H3K14ac was enriched in the Wnt-proximal cell, which also exhibited higher RNApolS2P, indicating enhanced transcriptional activity (Figure 2A-C).

**Figure 2:**
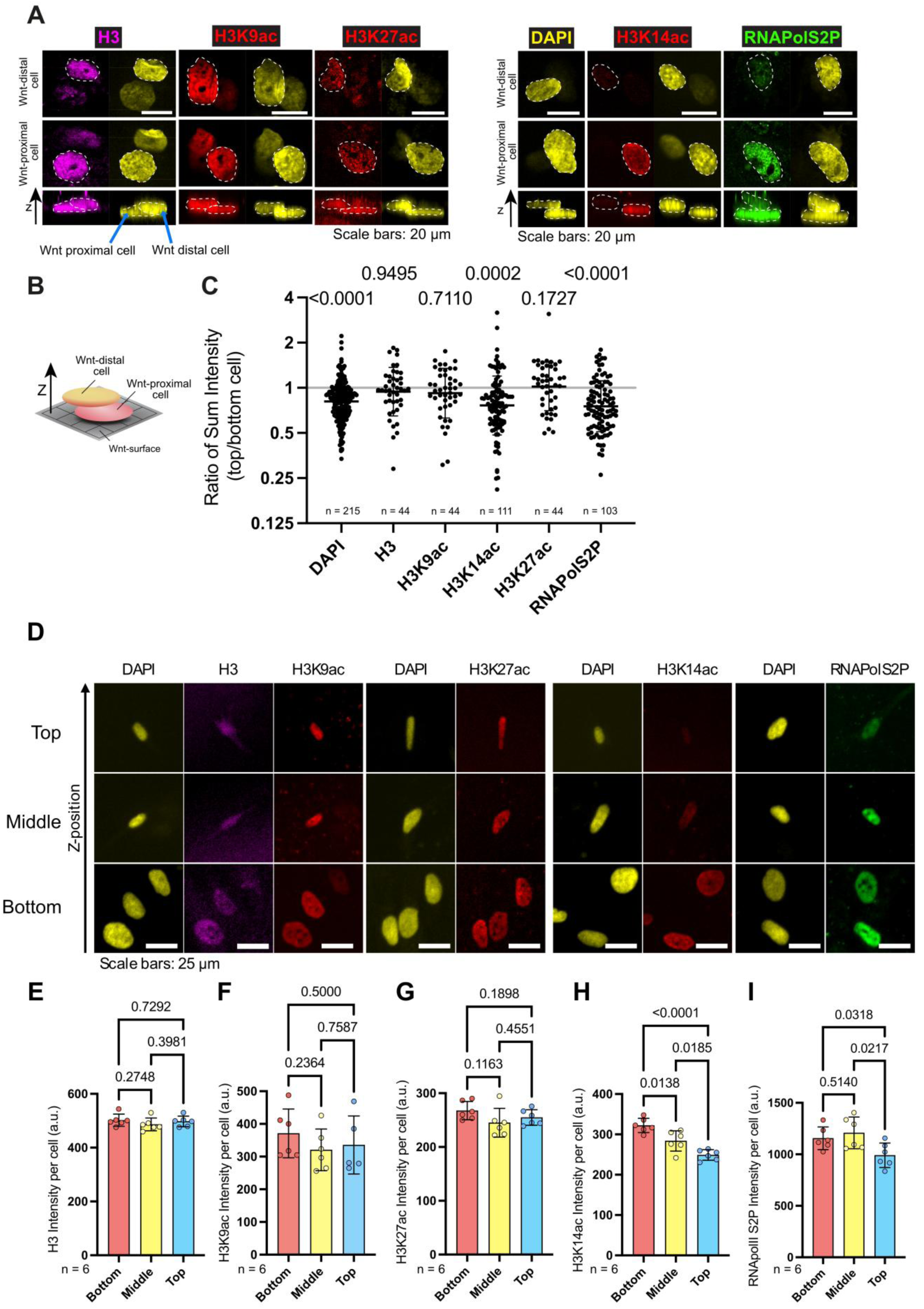
Asymmetric distribution of H3K14ac during hSSC asymmetric cell division and migration in the WIOTM. (A) Representative top- and side-view confocal images of hSSC doublets undergoing asymmetric cell division after 48 h of culture within the WIOTM. Cells were immunostained for DAPI (yellow), total histone H3 (magenta), acetylated histone marks H3K9ac, H3K14ac, H3K27ac (red), and phosphorylated RNA polymerase II (RNA Pol II S2P; green). (B) Schematic illustration of an asymmetric cell division oriented perpendicular to the Wnt3a platform in the WIOTM. (C) Quantification of the ratio of total fluorescence intensity between top and bottom daughter cells during asymmetric cell division, for each marker shown in (A). n represents the number of cell doublets analyzed. Values represent mean ± standard deviation (s.d.); statistical comparison was performed using a one-sample t-test against a theoretical ratio of 1 (equal distribution between cells). (D) Representative images of hSSCs localized in the Base, Middle, and Top layers of the WIOTM stained for the same epigenetic and transcriptional markers shown in (A). (E–I) Quantification of mean fluorescence intensity per cell for each marker within the three WIOTM layers: (E) H3, (F) H3K9ac, (G) H3K27ac, (H) H3K14ac, and (I) RNA Pol II S2P. n represents the number of WIOTM analyzed (≥ 95 cells analyzed per WIOTM). Data represent mean ± s.d.; statistical significance assessed by unpaired t-test with Welch’s correction.

Next, we examined H3 acetylation patterns in fully formed WIOTM by immunofluorescence. H3, H3K9ac, and H3K27ac showed no significant differences across Base, Middle, and Top regions. By contrast, H3K14ac was enriched at the Base, where hSSCs contact immobilized Wnt3a, and significantly reduced in the Middle and Top regions. RNApolS2P displayed a similar spatial trend, with comparable levels in the Base and Middle regions and a significant reduction in the Top region. Together, these data suggest an association between elevated H3K14ac and transcriptional activity near the Wnt3a interface (Figure 2D–I).

H3K14ac exists in a dynamic equilibrium maintained by histone acetyltransferases (HATs) and deacetylases (HDACs)^23^. Localized Wnt signaling increases H3K14ac, likely through enhanced HAT activity, reduced HDAC activity, or both, supporting sustained transcription and stem cell identity. We next wanted to know how this equilibrium is controlled.

### HDAC inhibitor SAHA abrogates Wnt3a-mediated asymmetric cell division and WIOTM formation

To investigate the role of H3K14ac in asymmetric cell division and WIOTM formation, we inhibited histone deacetylase (HDAC) activity using suberoylanilide hydroxamic acid (SAHA), a broad-spectrum HDAC inhibitor^29^. While SAHA is not specific for H3K14ac, given its inhibition of multiple HDACs that act on diverse histone modifications and non-histone proteins, it provides a means to assess the contribution of HDAC activity and histone acetylation to Wnt-mediated chromatin regulation.

Upon Wnt3a-mediated asymmetric cell division of hSSCs, daughter cells displayed differential levels of H3K14ac (Figure 3A–C), alongside variation in stem cell marker expression and osteogenic-bias marker expression^10^ (Figure 3B–D). In this study we used Cadherin-13 (CDH13) as a stem cell marker and Osteopontin (OPN) as an osteogenic-biased marker, as this pairing has been shown to faithfully recapitulate the behavior of the broader marker sets we and others previously established (STRO1, CDH13, PLXNA2 for stemness; OPN and OCN for osteogenic bias)^8,10,30^. Treatment with SAHA markedly reduced the differences in staining intensity of both DAPI and H3K14ac (Figure 3B–C) between daughter cells, whereas H3K9ac and RNAPol2S2P levels remained unaffected. SAHA treatment also abolished the asymmetry in osteogenic versus stem cell marker expression (Supplementary Table 1), an effect significantly different from the SAHA-solvent dimethyl sulfoxide (DMSO) control (Figure 3B, D).

**Figure 3:**
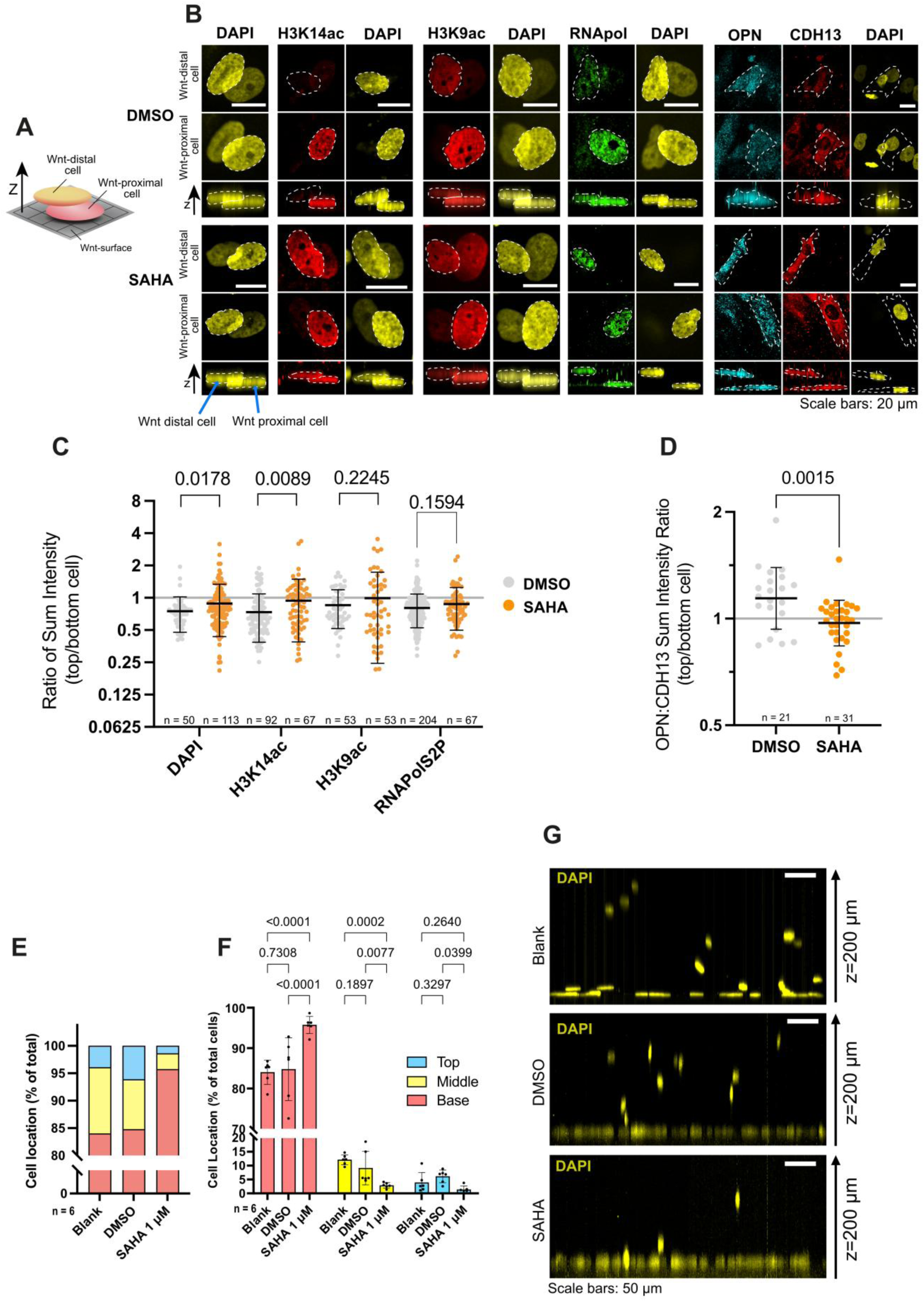
Inhibition of histone deacetylases alters asymmetric cell division of hSSCs, the asymmetric distribution of H3K14ac and WIOTM formation. (A) Schematic illustration of an asymmetric cell division oriented perpendicular to the Wnt3a platform in the WIOTM. (B) Representative top- and side-view images of hSSC doublets undergoing asymmetric division after 48 h of culture in the presence of either DMSO (vehicle control) or SAHA (1 µM). Cells were stained for DAPI (yellow), H3K9ac and H3K14ac (red), RNA polymerase II S2P (green), OPN (cyan), and CDH13 (red). Quantification of the ratio of total fluorescence intensity between the top and bottom daughter cells for markers in (B), comparing DMSO and SAHA conditions. (D) Quantification of the OPN:CDH13 fluorescence intensity ratio between the top and bottom cells during asymmetric division under DMSO or SAHA treatment. (C-D) n represents the number of cell doublets analysed. The data represents the mean ± s.d. Statistical analysis: ANOVA followed by unpaired t with Welch’s correction and one-sample t-test against a theoretical ratio of 1 (Supplementary Table 1). n ≥ 21 cell doublets. (E-F) Quantification of hSSC distribution across WIOTM layers (Base, Middle, Top) after 7 days of culture under three conditions: osteogenic media alone (Blank), osteogenic media + DMSO, and osteogenic media + SAHA (1 µM). The data represents the mean ± s.d. Statistical analysis: Uncorrected Fisher’s LSD tests. n = 6 WIOTM (≥ 109 cells per WIOTM). (G) Representative side-view projection images of hSSC migration in the WIOTM corresponding to the quantification in (E-F).

These findings indicate that SAHA disrupts the chromatin-based polarization induced by localized Wnt3a, abolishing fate asymmetry between daughter cells, even though cells continue to orient their spindles toward the Wnt3a source. This decoupling of spindle orientation and cell fate has also been observed in Wnt-induced asymmetric cell division of mouse embryonic stem cells^7^.

Since asymmetric cell division is required for WIOTM formation, we next examined whether SAHA similarly impaired WIOTM formation. After seven days in osteogenic medium containing 1 µM SAHA, cell migration within collagen gels was significantly altered: fewer cells were present in the middle and upper layers of the WIOTM, and more remained in the basal layer, compared with DMSO-treated and blank controls (Figure 3 E-G). Importantly, SAHA did not induce apoptosis in WIOTM cells, as assessed by TUNEL staining and Caspase-3 measurements (Supplementary Figure 1).

In summary, SAHA treatment leads to a broad-spectrum HDAC inhibition, which induces a general increase in acetylation, impairing Wnt-mediated asymmetric cell division of hSSCs and WIOTM formation.

### KAT7 downregulation abrogates Wnt3a-mediated asymmetric cell division and WIOTM formation

To specifically assess the role of H3K14ac in Wnt-mediated asymmetric cell division of hSSCs and WIOTM formation, we targeted KAT7, the principal histone acetyltransferase responsible for H3K14ac^31^. We downregulated KAT7 expression in hSSCs using a doxycycline-inducible shRNA system. We first verified KAT7 knockdown at the protein level by immunocytochemistry (Supplementary Figure 2A–B). Knockdown of KAT7 specifically reduced H3K14ac in transfected hSSCs without affecting a broad range of H4 acetylation (Supplementary Figure 2C) or other H3 acetylation marks (Supplementary Figure 3A).

During asymmetric cell division of shRNA-transduced hSSCs, doxycycline-induced KAT7 downregulation significantly reduced the asymmetric distribution of H3K14ac compared with both control shRNA and non-induced controls (Figure 4A–B). The asymmetry of osteogenic versus stem cell marker expression was also diminished, indicating disruption of Wnt-mediated asymmetric cell division of hSSCs (Figure 4C-D; Supplementary Table 2).

**Figure 4:**
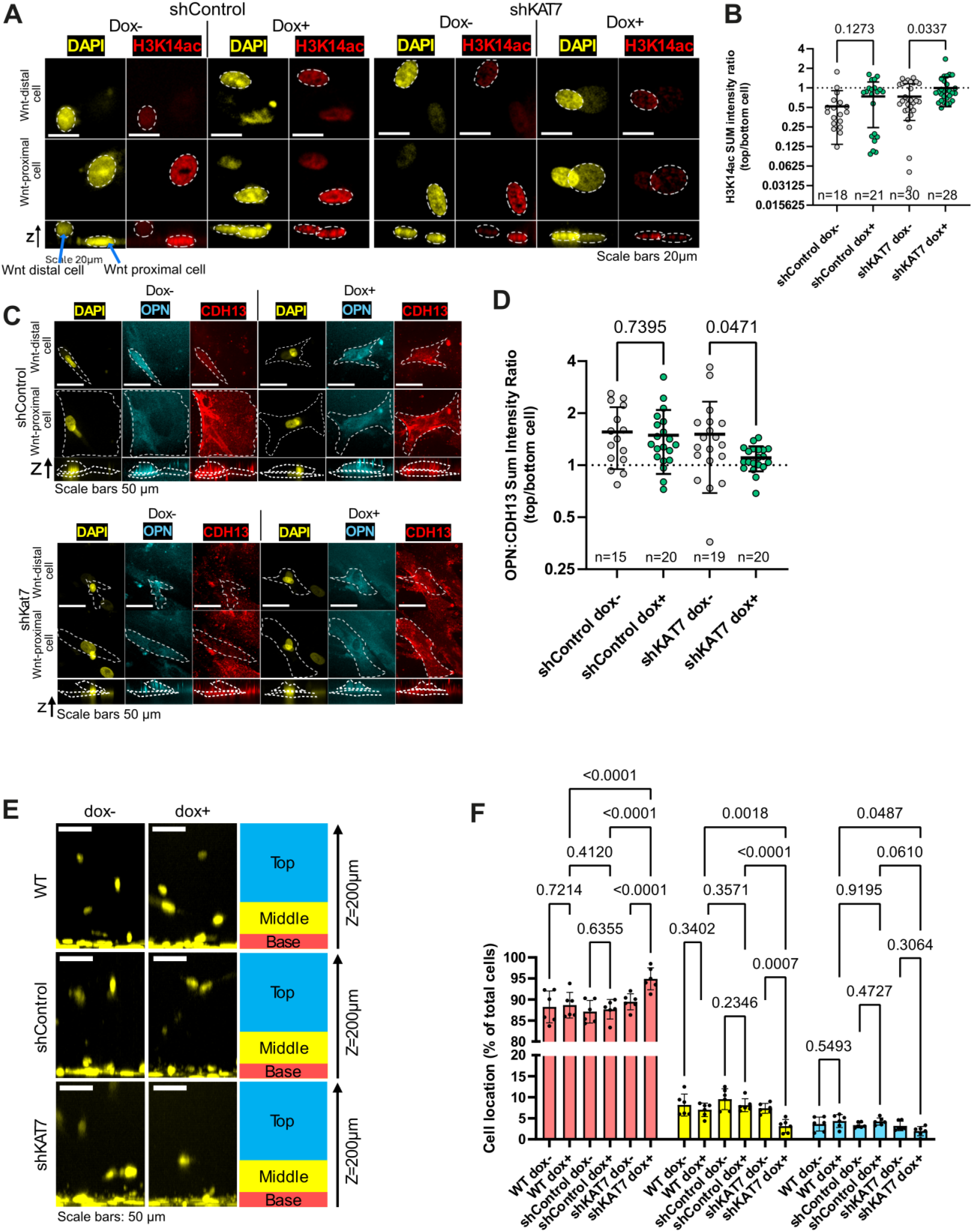
Kat7 knockdown disrupts H3K14ac and differentiation markers asymmetry during asymmetric cell division and alters migration of hSSCs in the WIOTM. (A) Representative top- and side-view images of hSSCs undergoing asymmetric cell division at the Wnt3a-platform after 48h of culture in the WIOTM. Cells were transduced with doxycycline (Dox)-inducible shRNA constructs targeting Kat7 or a non-targeting shRNA control. Staining was performed for DAPI and H3K14ac. (B) Quantification of H3K14ac ratio of total fluorescence intensity between the top and bottom daughter cells in the presence (green) or absence (grey) of doxycycline. (C) Representative top- and side-view images of hSSCs undergoing asymmetric cell division at the Wnt3a-platform after 48 h of culture in the WIOTM. Cells were transduced with doxycycline-inducible shRNA constructs targeting Kat7 or a non-targeting shRNA control. Staining was performed for DAPI, OPN and CDH13. (D) Quantification of OPN:CDH13 ratio between top and bottom cells under the same conditions as (B). Data in (B, D) are presented as mean ± s.d.. Statistical analysis: ANOVA followed by unpaired t with Welch’s correction and one-sample t-test against a theoretical ratio of 1 (Supplementary Table 2). Sample sizes: (B) n ≥ 18 cell doublets; (D) n ≥ 15 cell doublets. (E) Representative side-view projections of hSSCs migrating through the WIOTM after 7 days of culture. Comparisons include wild-type (WT) hSSCs, hSSCs transduced with Dox-inducible control vector, and hSSCs transduced with shRNA targeting Kat7, all cultured with or without doxycycline. (F) Quantification of cell distribution across WIOTM layers (Base, Middle, Top) after 7 days of culture for the conditions shown in (E). Data in (F) are presented as mean ± s.d. Statistical analysis was performed using uncorrected Fisher’s Least Significant Difference (LSD) test. Sample sizes: N = 6 WIOTM, n ≥ 94 cells.

Given that asymmetric cell division is a driver for WIOTM formation, we next examined whether KAT7 knockdown also disrupted this process. After seven days in osteogenic medium, doxycycline-induced KAT7 downregulation led to fewer cells in the upper layers of the WIOTM and increased retention of cells in the basal layer, relative to controls (Figure 4E–F).

Together, these results show that KAT7 depletion in hSSCs decreases H3K14ac, disrupts Wnt-mediated asymmetric hSSC division, and impairs WIOTM formation.

### Localized Wnt3a induces coordinated genome-wide chromatin remodeling in hSSC

To determine how immobilized Wnt3a globally reshapes chromatin accessibility, transcriptional output, and histone modification landscapes in hSSCs, we next performed genome-wide ATAC-seq, RNA-seq, and CUT&Tag profiling in hSSCs cultured on immobilized Wnt3a or inactive Wnt platforms.

Global chromatin accessibility profiles assessed by ATAC-seq differed between hSSCs cultured on immobilized Wnt and inactive Wnt platforms, with peak distributions showing moderate shifts in genomic localization (*p* value= 2.22e^-16^; Figure 5A) and a pronounced reduction in peak enrichment under immobilized Wnt conditions (Figure 5B). These changes are consistent with a global compaction of open chromatin regions following Wnt activation.

**Figure 5:**
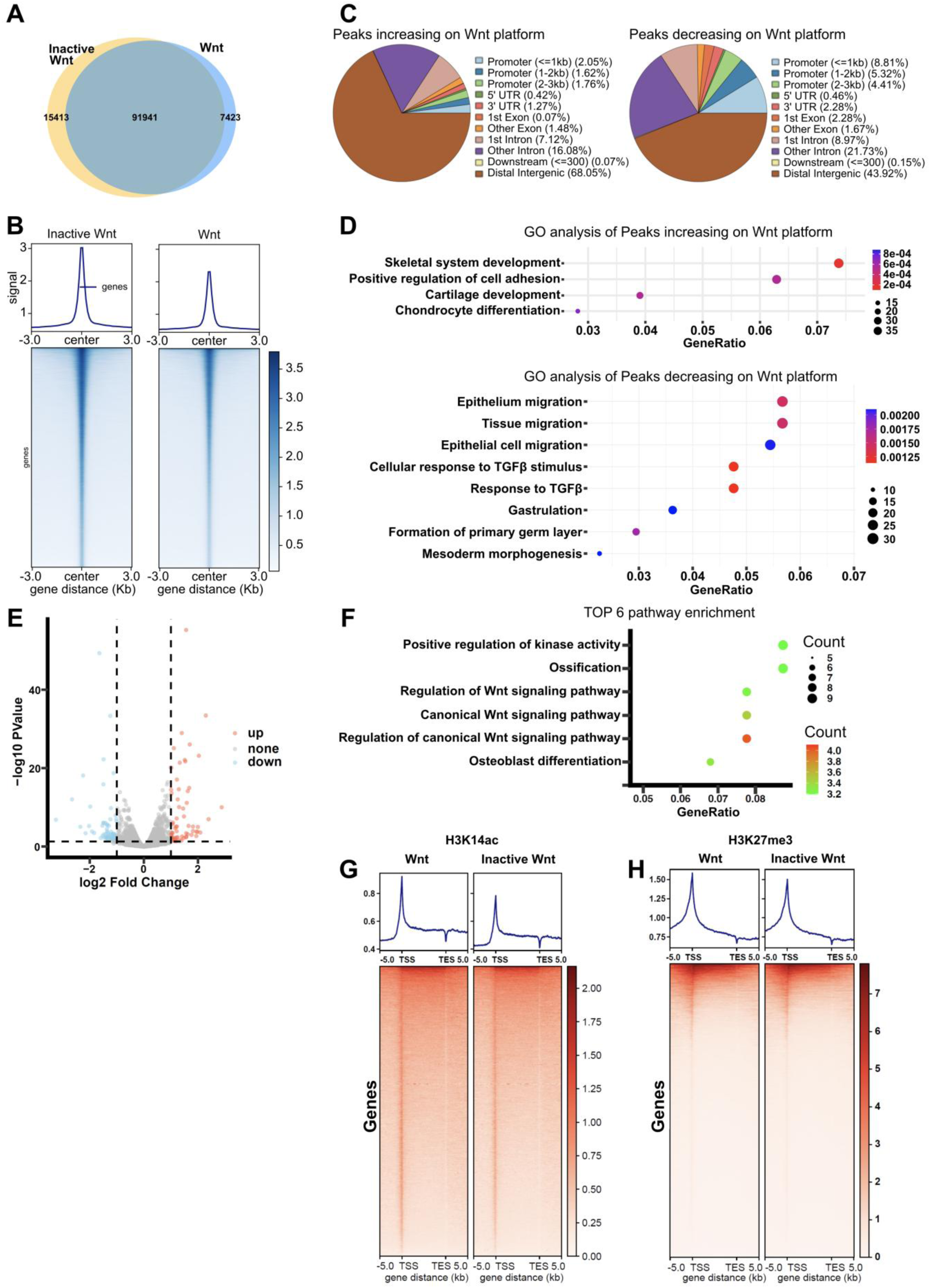
Activation of the Wnt signaling pathway increases H3K14ac levels in hSSCs. (A) Venn diagram showing the overlapping ATAC-seq peaks of hSSCs cultured on Wnt and DTT (dithiothreitol) inactivated Wnt. (B) The heatmap showing the enrichment of ATAC-seq signals 3000bp around the peak center. (C) Pie plots showing the distribution of significantly changed peaks on the genome after culture on Wnt. n = 1419 (Increased peaks), n = 661 (decreased peaks). (D) Dot plots showing the top 10 GO terms with the smallest P value. Dot color indicates the P value, and dot size indicates the enriched gene number. P values were calculated by Student’s t-test, two-sided. (E) Volcano plot showing differentially expressed genes (DEGs) between Wnt and inactive Wnt groups. Red points indicate up-regulated genes, and blue ones indicate down-regulated genes. Genes without any significant differences are in grey. P values were calculated by Student’s t-test, two-sided. n = 75 (down), n = 67 (up). (F) Dot plot showing the top 10 GO terms with the smallest P value for differentially expressed genes. Dot color indicates the P value, and dot size indicates the enriched gene number. P values were calculated by Student’s t-test, two-sided. (G) Profiles (top) and heatmaps (bottom) showing normalized H3K14ac signals on hSSCs cultured on Inactive Wnt and Wnt. 5 kb windows spanning the TSS (Transcription Starting Site) to TES (Transcription End Site) of all genes determined by UCSC Genes 2013 were plotted. Heatmaps were sorted by the average H3K14ac enrichment. (H) Same as G, except normalized H3K27me3 signals were used.

Although peaks with increased accessibility were fewer, they showed a distinct genomic pattern: they were depleted at promoters but enriched in intergenic regions (Figure 5C). Of the 1418 peaks with increased accessibility, 965 mapped to intergenic regions, 125 of which overlapped with known enhancers. Genes associated with these enhancer-overlapping peaks were frequently involved in skeletal system development, including osteogenic differentiation. Consistently, Gene Ontology (GO) analysis revealed that regions with increased accessibility were enriched for skeletal development and osteogenic differentiation pathways, whereas regions with decreased accessibility were associated with processes such as cell migration (Figure 5D, Sequencing Table 1 and 2).

To further examine the transcriptional consequences of Wnt activation, we performed RNA-seq on hSSCs cultured on Wnt or Inactive Wnt platforms. We identified 67 up-regulated and 75 down-regulated genes in Wnt–treated hSSCs (Figure 5E). Gene Ontology (GO) analysis revealed significant enrichment in the regulation of Wnt signaling and ossification (Figure 5F and Sequencing Table 3). To further investigate how Wnt signaling reshapes epigenetic states, we conducted CUT&Tag on hSSCs cultured on Wnt or Inactive Wnt platforms. When hSSc were cultured on Wnt platforms, a pronounced increase in H3K14ac enrichment was observed at promoter regions (Figure 5G). In contrast, H3K27me^3^ deposition showed only a minimal change (Figure 5H, Sequencing Table 5). Notably, genes showing Wnt-induced H3K14ac gains included multiple microRNA loci, such as MIR218-1, MIR34A, and MIR98 (Sequencing Table 4). These miRNAs are well-characterized regulators of Wnt signaling with stem cell fate regulation and osteogenic differentiation.

Among them, miR-218-1 has been shown to act as a feed-forward enhancer of the Wnt/β-catenin pathway by targeting inhibitors such as SOST, SFRP2 and DKK2, thereby amplifying Wnt signaling and driving osteoblast lineage commitment^32,33^. Similarly, miR-34A modulates osteoblast maturation and mineralization through repression of DKK1 and other osteogenic regulators^34^. In addition, miR-98-5p regulates osteoblast differentiation in MC3T3E1 cells by targeting CKIP1^35^.

Collectively, these results demonstrate that activation of the Wnt pathway induces coordinated remodeling of chromatin accessibility, transcription, and H3K14ac in hSSCs, highlighting the pivotal role of Wnt signaling in maintaining a transcriptionally active state during osteoblast differentiation and osteomimicry.

### H3K14-acetylation is maintained in transplanted hSSCs, delivered on various Wnt-biomaterials, in treating critical-sized calvarial bone defect

hSSCs are promising for tissue repair and are often delivered using biomaterial scaffolds that can influence their epigenetic state and cell fate^36–38^. Moreover, the *in vivo* environment presents a far more complex and dynamic setting than controlled *in vitro* culture, potentially altering epigenetic regulation in distinct ways. We therefore tested whether H3K14ac patterns are maintained in hSSCs and their progeny after WIOTM transplantation *in vivo*.

Our previous work demonstrated that a polycaprolactone (PCL)-based Wnt3a bandage promotes WIOTM formation, supports tissue survival *in vivo* for at least eight weeks, and ultimately contributes to bone repair^10^. In those experiments, we used hSSCs expressing mCherry, enabling lineage tracing of the transplanted cells and their progeny, and confirmed their identity by co-immunostaining with hSSC markers. Our results showed that hSSCs remained in close proximity to the bandage, whereas human cells migrating away downregulated hSSC markers and contributed to mature SOST⁺ bone cells within the mineralized bone tissue^10^.

Building on these findings, we re-analyzed the same samples to assess H3K14ac levels in human cells, identified by human β2-microglobulin (hB2M) marker. Human cells located adjacent to the PCL bandage or within the soft tissue near the bone exhibited higher H3K14ac levels compared with osteogenically differentiated human cells embedded within newly formed bone (Figure 6A–C). In contrast, human cells displayed comparable H3K9ac and H3K27ac levels regardless of their spatial proximity to the bandage (Supplementary figure 5A-D), mirroring what we observed in the WIOTM *in vitro* system (Figure 3B-C).

**Figure 6:**
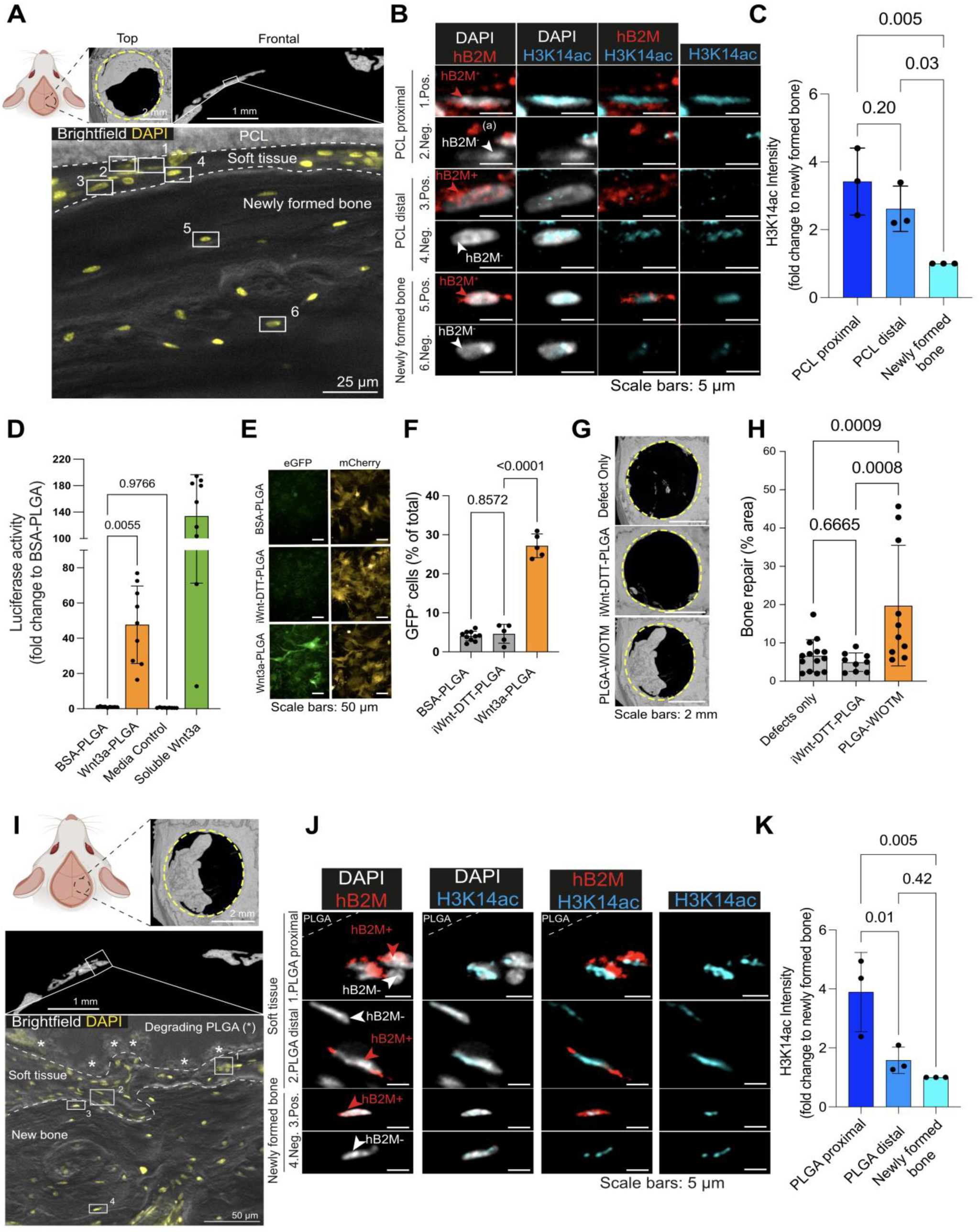
Localized Wnt-3a signaling on biodegradable polymers promotes bone repair, maintains and polarizes H3K14ac in transplanted hSSCs in a critical size defect model in mice calvaria. (A) Top panel: Schematic illustration of the calvarial defect surgical site, representative μCT images of calvarial defects with top view (Top), and digital frontal section (Frontal). In the Frontal view, the white square indicates the enlarged area. Bottom panel: representative image of the newly formed bone overlaid by the soft tissue, and the PCL bandage. The dashed line indicates the limit of each region. The white squares indicate the cell displayed in panel (B). (B) Representative images of cells of each tissue class: PCL proximal (i.e., close to the PCL), PCL distal (i.e., at distance from the PCL, close to the newly formed bone); newly formed bone (i.e., within the newly formed bone tissue). The numbers 1-6 refer to the position of the cell in the image of the panel (A). The human marker (human beta-2 microglobulin) is displayed in red and indicates cells of human origin, based on tissue-specific threshold (native bone). H3K14ac is displayed in blue with the same threshold in all images. (a) indicates the presence of staining artefacts (non-cellular origin). The red and white arrowheads indicate positive (hB2M+) and negative (hB2M-) cells, respectively. (C) H3K14ac intensity in cells of human origin in each tissue class. N = 3 defects (1 defect per animal), n = 296 cells. (D) Activity of immobilized Wnt3a on PLGA films measured by the 7xTCF-luciferase reporter cell line assay. N=9. (E) Wnt3a-induced eGFP expression of 7xTCF-eGFP//SV40-mCherry hSSCs on the PLGA bandage. mCherry is constitutively expressed in these reporter cells. BSA-PLGA and iWnt-DTT-PLGA bandages were used as negative controls. (F) Quantification of the Wnt3a-induced eGFP expression in (E). N³5. (G) Top views of representative μCT images of calvarial defects left uncovered (Defect Only), covered with iWnt-DTT-PLGA bandage, or covered with PLGA-WIOTM bandage. The dashed yellow circle indicates the edge of the drilled defects. (H) Quantification of the new bone area coverage over the defect site, as imaged by μCT. N³9 defects (one defect per animal). (I) Top panel: Schematic illustration of the calvarial defect surgical site, representative μCT images of calvarial defects with top view, and digital frontal section. The white square indicates the enlarged area. Bottom panel: representative image of the newly formed bone overlaid by the soft tissue, and the PLGA bandage. The dashed line indicates the limit of each region. The white squares indicate the cell displayed in the panel (J). (J) Representative images of cells of each tissue class: PLGA proximal (i.e., close to the PLGA), PLGA distal (i.e., at distance from the PLGA, close to the newly formed bone); newly formed bone (i.e., within the newly formed bone tissue). The numbers 1-4 refer to the position of the cell in the image of the panel (I). The human marker (human beta-2 microglobulin) is displayed in red and indicates cells of human origin, based on tissue-specific threshold. The red and white arrowheads indicate positive (hB2M+) and negative (hB2M-) cells, respectively. H3K14ac is displayed in blue with the same threshold in all images. (K) H3K14ac intensity in cells of human origin in each tissue class. N=3 defects (1 defect per animal), n = 194 cells. All data are presented as mean ± s.d.. Statistical analyses: ANOVA followed by uncorrected Fisher’s LSD.

Since PCL biodegrades over several years^39^, we sought to explore alternative Wnt-functionalized polymers with distinct degradation and mechanical characteristics for different repair settings. Polylactic-co-glycolic acid (PLGA), a clinically used resorbable polymer in applications such as sutures and some temporary fixation devices, typically degrades over 1–6 months depending on its lactide:glycolide ratio and formulation, thereby providing a faster-resorbing alternative^40^. Dynamic mechanical analysis at 37 °C revealed that PCL and PLGA bandages exhibit moduli of ∼350 MPa and ∼20 MPa, respectively (Supplementary Figure 6A-B). Although these values differ by an order of magnitude, both lie well above the stiffness range over which cells typically show strong sensitivity (1–100 kPa)^41,42^. Thus, at the cell–substrate scale, hSSCs are unlikely to directly sense this difference in bulk stiffness. In contrast, their divergent degradation kinetics, and the acidic byproducts associated with PLGA hydrolysis, represent biologically meaningful differences that could influence the repair microenvironment. We therefore investigated whether Wnt-PLGA-based materials could also be used to deliver WIOTM into critical-sized bone defects, thereby promoting repair and allowing analysis of H3K14ac patterns in human cells.

The immobilized Wnt3a onto PLGA retained biological activity, as shown by robust Wnt/β-catenin pathway activation *in vitro*. On PLGA–Wnt3a bandages, TCF/LEF-reporter LS/L cells^9,10^ exhibited a 47.7-fold increase in luciferase expression compared with inert BSA–PLGA controls (Figure 6D). Reporter hSSCs cultured on PLGA–Wnt3a bandages showed a five-fold increase in GFP^+^ cells (27.2%) relative to controls (∼5%) (Figure 6E-F). Given that only 53% of this population of hSSCs responds to Wnt3a^9^, the PLGA Wnt3a-bandage is capable of activating more than half of the responsive cells using this assay.

To evaluate the ability of PLGA to deliver the WIOTM in vivo and promote bone repair, we created 4 mm critical-size calvarial defects in 13-week-old immunocompromised female mice and transplanted PLGA–WIOTM bandages into the defect sites, following the same procedure used previously for PCL–WIOTM bandages^10^. We focused on female mice because their delayed healing and higher fracture risk make them a stringent model for bone repair^43,44^.

Similar to the first-generation PCL constructs, the PLGA–WIOTM bandage significantly enhanced bone regeneration compared with inert controls, achieving a mean repair of 19.8% in 4 mm defects (Figure 6G–H). Immunohistochemistry confirmed that the regenerated regions consisted of well-organized lamellar bone containing blood vessels (Supplementary Figure 4A).

Human-specific immunofluorescence staining (hB2M) revealed that within the defect site, 48.6% of cells located near the PLGA bandage (PLGA-proximal) were from human origin, 14% in soft tissue distant from the bandage (PLGA-distal), and 6.6% within the mature bone tissue (Supplementary Figure 4B–D). These findings demonstrate that the PLGA–WIOTM bandage supports both the survival and osteogenic differentiation of implanted human cells in vivo. As with the PCL–WIOTM bandage, human cells in proximity to the PLGA surface exhibited higher H3K14ac levels than human cells embedded in newly formed bone (Figure 6I-K). In contrast, human cells displayed comparable H3K9ac and H3K27ac levels regardless of their spatial proximity to the bandage (Supplementary figure 5A-D), mirroring what we observed in our PCL-WIOTM *in vivo* analysis (Figure 6A–C) and the WIOTM *in vitro* system (Figure 3B-C).

In summary, our results establish H3K14ac as a key epigenetic mark and regulator of hSSC fate during Wnt3a-mediated osteogenesis. In the 3D WIOTM model, nuclear morphology, chromatin condensation, and H3K14ac levels are spatially coordinated with stemness and differentiation, with H3K14ac specifically enriched in Wnt-proximal hSSCs. Disruption of H3K14ac, through HDAC inhibition or KAT7 knockdown, abrogates asymmetric cell division and impairs WIOTM formation, highlighting its critical role in orchestrating stem cell polarity and tissue patterning. Importantly, Wnt-functionalized PCL and PLGA bandages effectively deliver WIOTM *in vivo*, where they contribute to bone repair. Moreover, H3K14ac patterns are sustained in human cells within the osteogenic niche *in vivo*, mirroring those observed in WIOTM. Together, these findings reveal that targeted modulation of H3K14ac via Wnt-biomaterials provides a functional, tunable strategy to control stem cell fate and advance regenerative bone tissue engineering.

## Discussion

Regenerative medicine and tissue engineering require strategies that preserve stem cell populations while directing their differentiation into functional tissues. This balance is especially crucial in bone repair, where stem cell depletion compromises long-term regeneration^45^. Epigenetic mechanisms play a central role in maintaining this balance, integrating extrinsic signals with transcriptional programs that specify stem cell fate^46^. Using our Wnt-induced osteogenic tissue model (WIOTM), a 3D engineered construct that recapitulates aspects of the spatial and signaling complexity of the osteogenic niche, we link this control of hSSC behavior to an epigenetic regulator centered on H3K14ac.

In *Drosophila* germ cells^47^ and mouse embryonic stem cells^48^, it has been shown that certain histones and specific post-translational modifications are differentially distributed during asymmetric stem cell division, thereby influencing cell identity. Extending this to human primary stem cells, we find that H3K14ac is polarized between hSSC daughter cells of a Wnt3a-mediated asymmetric cell division, whereas H3K9ac, despite its frequent co-occurrence with H3K14ac at genomic loci^49,50^, does not show such distribution. Pharmacological inhibition of HDACs or genetic downregulation of KAT7, both of which alter H3K14ac levels, disrupt Wnt-mediated asymmetric hSSC division and osteogenic differentiation, supporting a model in which H3K14ac acts not merely as a marker but as a determinant of lineage outcome. This functional role likely reflects divergent regulation by specific lysine acetyltransferases, including KAT7, and the broader activity spectra of histone deacetylases^51^.

Genome-wide profiling revealed that localized Wnt signaling drives extensive chromatin remodeling. ATAC-seq showed a global reduction in chromatin accessibility, consistent with tighter chromatin packing, accompanied by a selective opening of enhancer-associated intergenic regions enriched for skeletal developmental pathways. Regions that closed under Wnt activation were instead associated with cell migration and early developmental programs, suggesting that Wnt restricts hSSCs from alternative trajectories while reinforcing osteogenic commitment. Transcriptomic profiling corroborated these findings through the induction of canonical Wnt targets and osteogenic transcription factors. Concurrently, CUT&Tag demonstrated a pronounced increase in promoter-associated H3K14ac, including at loci linked to MiR-218-1 modulators such as SFRP2 and DKK2, extracellular regulators of Wnt ligand and Wnt signaling^52^, while H3K27me3 remained largely unchanged. These findings indicate that Wnt signaling promotes a transcriptionally permissive chromatin landscape primarily through enhanced histone acetylation rather than altered repressive marking. Collectively, these genome-wide data demonstrate that localized Wnt activation reconfigures the epigenome by selectively enhancing acetylation and accessibility at osteogenic regulatory elements, thereby reinforcing transcriptional programs essential for skeletal lineage commitment.

Mechanical cues also influence osteogenic differentiation^36^, often acting through HDAC-dependent pathways that modify histone acetylation^53,54^, including H3K14ac^55^. These biomechanical effects underscore the need to study human osteogenesis in 3D contexts that reflect the spatial organization of bone rather than relying solely on 2D systems^56^. Indeed, 2D osteogenic models frequently identify H3K14ac as a pro-osteogenic mark, promoting genes such as *Runx2*^57,58^. Similarly, osteogenic differentiation of osteoblasts in 2D is promoted by SAHA inhibition of HDACs, resulting in H3K14 hyperacetylations^59^. Yet *in vivo*, SAHA treatment leads to bone loss by inhibiting immature osteoblasts^60,61^, highlighting discrepancies between 2D and physiological environments. The WIOTM resolves this gap: in 3D, HDAC inhibition disrupts Wnt-mediated asymmetric hSSC division and osteogenesis, revealing the context-dependent roles of histone modifications and the limitations of 2D models for predicting regenerative outcomes.

The translational utility of the WIOTM is further demonstrated by its incorporation into Wnt-functionalized bandages fabricated from either PCL^10^ or PLGA. Although these polymers differ in degradation kinetics, with PCL degrading slowly over years and PLGA degrading within months and releasing acidic byproducts, both materials supported WIOTM formation and comparable bone repair outcomes, indicating that local Wnt cues dominate over polymer chemistry in guiding hSSC behavior. This is particularly important given that conventional PLGA scaffolds, while widely used, often promote disorganized woven bone formation and release acidic degradation products that impair osteogenesis^62–65^. By contrast, the WIOTM structure spatially separates differentiating cells from the PLGA surface, enabling orderly osteogenic maturation as cells migrate away from the Wnt3a-functionalized interface. Strikingly, we find that human cells in direct proximity to either PCL- or PLGA-based Wnt bandages maintain high levels of H3K14ac, which become progressively downregulated as cells migrate away from the scaffold, mirroring the spatial pattern observed within the WIOTM itself. Although immobilized Wnt3a is unlikely to remain biochemically active eight weeks post-transplantation, this persistence of H3K14ac likely reflects the reinforcement by endogenous Wnt signals from the injury niche^3,66,67^. Such maintenance of epigenetic identity under physiologically heterogeneous conditions is critical, as it preserves the hSSC pool, prevents premature differentiation, and supports organized tissue regeneration rather than disordered repair. Collectively, these findings demonstrate that the WIOTM recapitulates not only transcriptional hallmarks of osteogenesis but also certain spatial organization of epigenetic states observed *in vivo*.

Together, our findings position H3K14ac as a mechanistic link between Wnt signaling and hSSC fate in controlled systems, while the *in vivo* observations support the relevance of this axis in regenerative settings. Looking forward, the WIOTM provides a powerful platform to investigate additional epigenetic modifications and their molecular mechanisms in both engineered tissues and transplanted grafts. Because the WIOTM bandage can be transplanted into bone defects, it enables direct interrogation of how stem cell epigenetics adapt to and persist within the complex *in vivo* repair environment. Moreover, the WIOTM can be employed as a drug-screening platform to identify epigenetic modifiers that enhance osteogenesis while preserving stemness, with potential direct applications to bone repair. Beyond regenerative medicine, this approach may also yield insight into pathological contexts such as cancer, where dysregulated asymmetric division and epigenetic control of stemness drive disease progression.

## Methods

### Wnt3a purification

Recombinant mouse Wnt3a was produced in Drosophila S2 cells grown in suspension culture. Wnt3a was purified by Blue Sepharose affinity and gel filtration chromatography, and biological Wnt3a activity was determined in a luciferase reporter assay using L cells stably transfected with the SuperTOPFlash reporter (LS/L), as described previously^8^. Alternatively, human recombinant Wnt3a protein (R&D Systems, 1324-WN-010) was purchased from commercial vendors and tested following the same procedures, before use.

### Wnt3a immobilization and validation

Wnt3a platforms were prepared on amine-coated wells (Corning PureCoat Amine 96-well flat-bottomed multiwell plates, VWR International, 734-1475), as previously described^8–10^. Wells were washed once with 100 µL Dulbecco’s PBS (dPBS) and then incubated with 50 µL of 5% (v/v) glutaraldehyde (Sigma-Aldrich, G6257) in H_2_O for 30 min at room temperature (RT) in the dark. The wells were washed three times with 100 µL dPBS, incubating each wash for 5 min at RT. Next, the glutaraldehyde-activated wells were incubated with 60 ng Wnt3a in dPBS for 1 h at room temperature, in the dark. After Wnt3a immobilization, wells were washed three times with 100 µL dPBS, incubating each wash for 10 min at RT. For the inactivated Wnt3a (iWnt3a) platforms, 100 µL of 20 mM DTT (Life Technologies, P2325) in PBS was added to the wells and incubated for 30 min at 37 °C. The wells were washed three times with 100 µL dPBS, incubating each wash for 10 min at RT. After removing the final wash, the wells were incubated with hSSC basal medium for at least 1 h before being used for downstream purposes. For each plate prepared, Wnt3a-immobilised wells were used to validate biological Wnt3a activity after immobilization. Briefly, the fluorescence intensity from Wnt-responsive COMMA-D beta cells stably infected with 7xTCF-eGFP// SV40-mCherry was measured after imaging with an epifluorescent microscope (Nikon). Before imaging, the cells were stained with Hoechst 33342 (ThermoFisher, H3570) for 5 min and then washed with dPBS.

### Human skeletal stem cell culture and expansion

hSSCs were purchased from Lonza (PT-2501). hSSCs were expanded up to passage 5 (P5) in ‘basal medium’: high glucose (4.5 g.L^−1^) Dulbecco’s modified Eagle’s medium (DMEM; Sigma-Aldrich, D6546) supplemented with 10% fetal bovine serum (FBS; Sigma-Aldrich, F7524) and 1% penicillin/streptomycin (Sigma-Aldrich, P4458) at 37°C in humidified air with 5% CO₂.

### Knockdown of Kat7 in hSSCs

hSSCs expressing an inducible shRNA targeting Kat7 were generated using lentiviral particles. Briefly, TRIPZ constructs targeting Kat7 (RHS4696-200710314) and control constructs (scramble, RHS4743; empty, RHS4750) were bought from Horizon Discovery as *E. coli* glycerol stocks. After bacterial culture overnight in 7 mL of LB broth (Sigma, L3522) + ampicillin 100 µg/mL (Sigma, A9518), and plasmid purification using the High Pure Plasmid Isolation Kit (Roche, 11754777001), constructs were transfected in HEK293T LentiX cells (Takara Bio, 632180) seeded in T75 flasks along with the Trans-Lentiviral Packaging Mix(Horizon Discovery, TLP5913) for lentiviral particle production according to the manufacturer’s recommendations. Lentiviral particles were collected after 48 h of production, concentrated using Lenti-X™ Concentrator (Takara Bio, 631231), and resuspended in 100 µL of sterile PBS. Then, 75000 hSSCs per well were seeded in a 6-well plate and cultured overnight before transduction with 30 µL of lentiviral stock. Four days after transduction, transfected cells were selected by adding 2 µg/mL of puromycin (ThermoFisher, J61278) to the culture media for 7 days, changing media every 2–3 days. hSSCs were further amplified using basal culture media. shRNA expression induction was performed by adding doxycycline (ThermoFisher, J60422.06) to the culture media at a final concentration of 2 µg/mL, changing media every day for 7 days.

### 3D culture of hSSCs to assess asymmetric cell division and WIOTM formation

2,500 hSSCs per well were seeded on the Wnt3a platform and allowed to attach for 24 h. Then, media was aspirated, and cells were overlaid with 100 µL/well collagen gel (0.5 µg/mL rat tail collagen I, Corning, 354249) diluted in high-glucose DMEM plus 3.3 µM NaOH and used immediately. Gelation was allowed for 10 min at RT and then for 2 h at 37 °C, 5% CO₂, before 150 µL osteogenic media was added to each well. Plates were returned to the incubator (37 °C, 5% CO₂) and cultured for 48 h (for ACD analysis) or 7 days (for WIOTM formation). Osteogenic media was gently aspirated and replaced every 2 to 3 days.

### hSSCs SAHA treatment

Suberoylanilide hydroxamic acid (SAHA, Sigma, SML0061) was dissolved in DMSO at a stock concentration of 1 mM and was added to the osteogenic media at a final concentration of 1 µM. Cells were cultured in the presence of the inhibitor for 48 h (ACD) or 7 days (WIOTM formation), replacing media every 2–3 days.

### TUNEL assay and staurosporine treatment

Cells were cultured for 7 days to allow WIOTM formation (see section above), in the presence or absence of SAHA (final concentration: 1 µM). Culture medium was replaced every 2–3 days. Six hours before fixation on day 7, three wells were treated with staurosporine (10 µM; Fisher Scientific, 15495139) to serve as a positive control for cell death. Cells were fixed in 4% paraformaldehyde (PFA) for 20 min at room temperature and subsequently processed for apoptosis detection using the Click-iT™ Plus TUNEL Assay Kit (ThermoFisher, C10619). Parallel samples were immunostained for cleaved caspase-3 as described in the section below.

### 3D immunostaining of asymmetric cell division and WIOTM

At the experimental endpoint, media was gently aspirated and the well rinsed with PBS, taking care not to disturb the collagen gel. Cells and gel were fixed with 200 µL 4% PFA for 10 minutes at RT. Fixative was removed and the well was washed twice with 200 µL PBS. Permeabilization was performed with 200 µL 0.3% Triton X, 0.05% Tween-20, 0.05% Sodium Azide in PBS, incubating for 3 h at RT before aspiration. Primary antibodies were diluted to the required concentration in antibody buffer (1% BSA, 0.1% Tween-20, 0.05% Sodium Azide in PBS) and added to the well for 72 h at 4 °C. Wells were washed four times for 30 minutes with PBST (0.1% Tween-20 in PBS). Secondary antibodies and DAPI were diluted in blocking buffer at 1:1000 and added to the well overnight (∼16 hours) at 4 °C. Wells were washed four times for 30 minutes with PBST (0.1% Tween-20 in PBS). 150 µL of storage solution (0.1% Sodium Azide in PBS) was added to the wells to allow imaging for up to 1 week and plates were stored at 4 °C.

### Antibodies

The antibodies used for immunofluorescence were: H3 (mouse, 1:250; Cell Signaling, 14269S), H3K9ac (Rabbit, 1:250; Abcam, ab10812), H3K14ac (Rabbit, 1:250; Abcam, ab52946), H3K18ac (Rabbit, 1:250; Invitrogen, PA-585523), H3K23ac (Rabbit, 1:250; Sigma, SAB5600023), H3K27ac (Rabbit, 1:250; Abcam, ab4729), H3K36ac (Rabbit, 1:250; ThermoFisher, MA5-24672), H4 (mouse, 1:250; ThermoFisher, MA550233), H4K5ac (Rabbit, 1:250; Invitrogen, SAB5600018), H4K8ac (Rabbit, 1:250; Sigma, SAB5600019), H4K12ac (Rabbit, 1:250; Abcam, ab46983), H4K16ac (Rabbit, 1:250; Invitrogen, 720083), RNAPolS2P (Rat, 1:250; Abcam, ab252855), Osteopontin (Mouse, 1:50; Santa Cruz, sc21742), Cadherin-13 (Rabbit, 1:50; Abcam, ab36905), KAT7 (Rabbit, 1:250; Abcam, ab70183), Cleaved Caspase 3 (Rabbit 1/250; Cell Signaling, 9579).

### Microscopy

Imaging after 3D immunostaining was performed using an inverted spinning disk confocal microscope (Nikon, NIS-Elements Viewer 5.2 software). ACDs were imaged using a 40x (NA = 0.95) objective, with a step size of 0.6 µm, determined according to the Nyquist criteria. WIOTMs were imaged using a 20x (NA = 0.75) objective, using a step size of 3 µm. Resulting confocal data were analyzed in ImageJ as described in the section below.

### Image analysis in ImageJ

#### WIOTM analysis

To analyze the intensity and distribution of DAPI-stained nuclei in the WIOTM, files were manually analyzed to locate each cell along the Z-axis. A square selection was drawn around the cell, the highest and lowest Z-positions encompassing the cell were chosen, and the “Duplicate” function was used to generate an independent 3D stack, which was then used for analysis. The average position of the cell (bottom Z-position + top Z-position) / 2 was used as the reference position of the cell in the gel. All the cells attached to the bottom were analyzed together. The “Stack > Z project… > Max projection” or “Stack > Z project… > Sum projection” function was used to generate a maximum or sum intensity projection of each stack. Then, a manual region of interest (ROI) was drawn around the nucleus periphery of each cell, and the “Measure” function was used to measure nucleus area, circularity, roundness, and fluorescence intensity (Mean or Sum intensity) for each channel in the image. Measured information for each cell, along with its position in the collagen gel, was recorded in an Excel file. To plot and analyse the data, cells were divided according to their position in the collagen gel as “Base” (cells in direct contact with the bottom of the plate), “Middle” (cells found within the first 75 µm of collagen gel but not in the “Base” category), and “Top” (cells found above 75 µm). Histone acetylation and RNApolS2P immunostaining in the WIOTM were analyzed using the “Stack > Z project… > Max projection” function to generate a maximum intensity projection of each WIOTM layer (Base, Middle, Top), according to the DAPI analysis. Then, a manual region of interest (ROI) was drawn around the nucleus periphery of each cell, and the “Measure” function was used to measure mean fluorescence intensity for each channel in the image.

### ACD analysis

Confocal images containing post-division doublets (identified as two cells closely overlaid in the z-axis) were used to analyze asymmetric histone marks and cell fate marker distribution during ACD.

For histone mark analysis, one Z-slice per cell was selected using predefined criteria of maximal nuclear signal intensity. Quantification was performed using a manually drawn ROI encompassing the nuclear boundary to measure mean fluorescence intensity and nuclear area.

Fluorescent intensity of the background was also measured. Data was transferred to an Excel file for post-analysis. Normalized fluorescent intensity was calculated by subtracting the background intensity and multiplying by the nucleus area, and was used to calculate the intensity ratio as: ratio = Norm. intensity Top cell / Norm. intensity Bottom cell. For presentation purposes, the log₂ of the ratio was calculated and plotted using GraphPad Prism.

For cell fate markers, a single “optimal” Z slice in which the staining intensity was brightest and most homogeneous throughout the cell was chosen for analysis for each cell (bottom and top). A manual ROI drawn around the cell periphery of each cell was used to measure mean fluorescence intensity for the bottom and top cell of each marker (OPN and CDH13). Fluorescence intensity of the background was also measured. Data were transferred to an Excel file for post-analysis, and normalized fluorescent intensity was calculated by subtracting the background intensity. For each cell, the ratio of cell fate markers was calculated for each cell as follows: Norm. intensity OPN / Norm. intensity CDH13. Then, the calculated top cell ratio was divided by the calculated bottom cell ratio as follows: Top cell Ratio OPN:CDH13 / Bottom cell Ratio OPN:CDH13. For presentation purposes, the log₂ of the ratio was calculated and plotted using GraphPad Prism.

### ATAC sequencing

hSSCs were seeded into 96 well plates at 3000 cells per well onto Wnt3a platform or Inactive Wnt3a platform with 100 µL hSSC basal medium and cultured for 24h to allow cells to attach. After 24h, media was replaced with osteogenic media and cells were cultured for 48h. Cells were washed twice with PBS and trypsinized (Trypsin EDTA 0.05%) for 3min at 37C. 18 wells were pooled to obtain enough cells and counted as 1 biological replicate (n=4 per group). Samples were then processed for ATAC-seq using a commercial kit (Active Motif, ATAC-Seq Kit) according to the manufacturer’s recommendation. Sample quality was assessed using the Fragment Analyzer 5200 (Agilent, software version: 3.1.0.12). Sequencing of the libraries generated was done on an Aviti sequencer (Element Biosciences), 50 million reads per library.

### ATAC analysis

To remove adaptors and low-quality reads, Trim Galore (version 0.6.4) (https://www.bioinformatics.babraham.ac.uk/projects/trim_galore) was used with the parameter ‘--paired’. Trimmed reads were then mapped to the human reference genome (hg19) using Bowtie2^68^ (version 2.3.5.1). PCR duplicates were removed by GATK4 (version 4.1.4.0) (https://github.com/broadinstitute/gatk) with the parameter ‘--REMOVE_DUPLICATES = true’. MACS2^69^ (version 2.2.6) was used to call narrow peaks for ATAC-seq. For visualizing epigenomic signals, normalizing mapped reads, and calculating the coverage of features across the genome, BEDtools^70^ (version 2.92.2) and the bedGraphToBigWig approach (version 4) (https://www.encodeproject.org/software/bedgraphtobigwig) were used. deepTools^71^ (version 3.4.3) was used to draw heatmaps by the function computeMatrix and plotHeatmap. For downstream differential peaks analysis, multiBigwigSummary in deepTools was used to count average signals over each region. Differential regions analysis was performed by limma^72^ (version 3.42.2) with normalized signals. clusterProfiler^73^ (version 3.14.3) was used for peak distribution analysis, peak annotation, and GO analysis. The known enhancer annotation was from the database EnhancerAtlas 2.0^74^.

### RNA sequencing

hSSCs were seeded into 96 well plates at 3000 cells per well onto Wnt3a platform or Inactive Wnt3a platform with 100 µL hSSC basal medium and cultured for 24 h to allow cells to attach. After 24 h, media was replaced with osteogenic media and cells were cultured for 48 h. RNA was extracted and purified using a commercial kit (Nucleospin RNA XS, Macherey-Nagel) according to the manufacturer’s recommendation. 30 wells were pooled to obtain enough material and counted as 1 biological replicate (n=3 per group). Samples were sequenced using the NovaSeq6000 instrument (Illumina).

### RNA analysis

To remove adaptors and low-quality reads, Trim Galore (version 0.6.7) (https://www.bioinformatics.babraham.ac.uk/projects/trim_galore) was used with the parameter ‘--paired’. STAR^75^ (version 2.7.6a) was used for mapping the filtered reads to the human reference genome (hg19). Expression counts were generated using featureCounts^76^ (version 2.0.1). Differentially expressed genes were identified by the R package DESeq2^77^ (version 1.44.0) with the criteria of "p.value < 0.05 and |log_2_FC| >= 1". clusterProfiler^73^ (version 4.12.3) was used to perform Gene Ontology (GO) analysis.

### Cut&Tag

hSSCs were seeded into 96 well plates at 3000 cells per well onto Wnt3a platform or Inactive Wnt3a platform with 100 µL hSSC basal medium and cultured for 24 h to allow cells to attach. After 24 h, media was replaced with osteogenic media and cells were cultured for 48 h. Cells were washed twice with PBS and trypsinized (Trypsin EDTA 0.05%) for 3 min at 37°C. 15 wells were pooled to obtain enough cells and counted as 1 biological replicate (n=3 per group). Samples were then processed for Cut&Tag using a commercial kit (Active Motif, CUT&Tag-IT Assay Kit Anti-Rabbit) according to the manufacturer’s recommendation. Antibodies used: anti-H3K14ac (Abcam, Ab52946), anti-H3K27me3 (Invitrogen, MA5-11198). Sample quality was assessed using the Fragment Analyzer 5200 (Agilent, software version: 3.1.0.12). Samples were sequenced using the NovaSeq6000 instrument (Illumina), 50 million reads per library.

### CUT&Tag analysis

To remove adaptors and low-quality reads, Trim Galore (version 0.6.7) (https://www.bioinformatics.babraham.ac.uk/projects/trim_galore/) was used with the parameter ‘--paired’. Then, trimmed reads were uniquely mapped to the human reference genome (hg19) using Bowtie2^68^ (version 2.3.5.1). The output SAM files were converted to sorted BAM files using SAMtools^78^ (version 1.9). For visualizing enrichment signals, normalizing mapped reads, and calculating the coverage of features across the genome, BEDtools^70^ (version 2.26.0) and bedGraphToBigWig (version 4) (https://www.encodeproject.org/software/bedgraphtobigwig/) were used with the following parameters ‘-scale 1e^7^/(the total number of mapped reads)’. deepTools^71^ (version 3.4.3) was used to draw heatmaps by the function computeMatrix and plotHeatmap.

### Manufacturing of biodegradable polymers

PCL (Mn = 80,000; Merck, 440744), or PLGA (Mn = 190,000–240,000; Merck 739979) sheets weighing 0.3 g were created using the solvent-casting method in 7-cm glass dishes by dissolving the polymer at a concentration of 2% (w/v) in anisole (Merck, 296295) and were then left for 72 hours at 50 °C to facilitate the evaporation of the solvent.

### Glutaraldehyde group introduction into biodegradable polymers

The desiccated PLGA films were meticulously detached from the dishes and treated with 2% (w/v) HMDA (Sigma-Aldrich, H11696) in 90% (v/v) isopropanol (Sigma-Aldrich, I9516) for 1.5 hours on a tube roller at ambient temperature to introduce amine groups onto their surface. Subsequently, the films were rinsed three times for 10 minutes each in 100% isopropanol and allowed to air dry.

The dried PCL films were plasma-activated using a plasma etching electrode in oxygen gas at a flow of 20 cm³ min⁻¹ with a radio frequency plasma generator (frequency 13.56 MHz, power 50 W, Diener electronic Zepto-W6) at 0.2 mbar pressure for 3 min. The freshly plasma-activated films were immediately treated with 5% APTES (Sigma-Aldrich, 440140) in ethanol (Sigma-Aldrich, 32221) for 2 h. The films were then washed in 100% ethanol three times for 10 min each and left to dry. The PLGA and PCL films were further treated with 5% (w/v) glutaraldehyde (Sigma-Aldrich, G6257) in 70% (v/v) ethanol for 5 min at room temperature to introduce aldehyde groups onto the surface of the polymer. The polymers were then washed three times with 100% ethanol and left to dry. The polymers were finally sterilized in 100% ethanol for 15 min before being washed three times in PBS.

### Wnt3a modification of biodegradable polymers

Wnt3a protein (RnD), was diluted in PBS for a final volume of 20 µL containing 300 ng of Wnt3a. The polymer films were incubated with soluble Wnt3a for 1 h, washed three times with dPBS, and then further incubated with high-glucose DMEM containing 20% FBS to block any unreacted aldehyde groups. iWnt3a films were also produced as controls. This was achieved by inactivating the Wnt3a-modified polymers with 20 µL of 20 mM DTT for 30 min at 37 °C.

### Culture of hSSCs on Wnt3a-functionalized biodegradable polymers

For *in vitro* analysis of the functionalized films, hSSCs with the 7xTCF-eGFP//SV40-mCherry reporter were seeded onto the functionalized PLGA films at 35,000 cells cm⁻² (10,000 cells per bandage) and cultured in hSSC basal medium for 24 h. For the preparation of the *in vivo* cellular bandages, the functionalized polymer films (Wnt3a in WIOTM-bandages, inactive Wnt3a in iWnt3a-bandages + hSSCs) were seeded with cells at the same density and allowed to adhere. The basal medium was then removed and 100 µL of 1 mg mL⁻¹ rat tail collagen I was overlaid, allowed to set, and topped up with osteogenic medium (details). WIOTM-bandages and iWnt3a-bandages + hSSCs were cultured for 7 days, respectively, with three media changes.

### Analysis of Wnt activity on Wnt3a-functionalized PLGA

The fluorescence intensity from Wnt-responsive hSSCs stably infected with 7xTCF-eGFP//SV40-mCherry was measured using an Operetta High-Content Imaging System (PerkinElmer). Before imaging, the cells were stained with Hoechst 33342 (ThermoFisher, H3570) for 5 min, washed with dPBS, and then the medium was refreshed with 20 µL dPBS containing 10% FBS. We measured the Wnt activity of functionalized biodegradable polymers by culturing the L cells transfected with SuperTOPFlash reporter (LS/L), as previously reported, in DMEM supplemented with 10% FBS and 1% penicillin/streptomycin overnight, and quantified the Wnt-induced luciferase activity using a Dual-Light System (Applied Biosystems).

### Transplantation of osteogenic and control bandages

Thirteen-week-old female SCID mice (Charles River, Strain 236) were housed in conditions that comply with the Animals (Scientific Procedures) Act 1986 (ASPA). All procedures were performed under sterile conditions. Firstly, general anaesthesia was administered to the mice by the intraperitoneal route using an anaesthetic cocktail (10 µL per g body weight) consisting of sterile sodium chloride 0.9% saline with 7.5 mg mL⁻¹ ketamine (Vetalar) and 0.1 mg.mL⁻¹ medetomidine (Domitor). After testing for the loss of tail and hind paw withdrawal reflex, the top of the head was shaved with a mini hair trimmer and lidocaine 5% w/w ointment applied to promote local anaesthesia. The eyes were lubricated with Viscotears liquid gel, and then a sagittal skin incision was made along the midline of the head with the scalpel. The mice were positioned under the stereoscopic microscope, the cranial vault was exposed, and the periosteal layer overlying the skull was removed with an outward separating motion using cotton buds. One 4 mm defect per animal (n≥ 9 animal per group) was drilled gently using a dental handpiece, avoiding the build-up of heat by friction. Removal of bone pieces was followed by covering the defects with 40 µL of 1 mg.mL⁻¹ rat tail collagen I, placing the bandage treatment onto the defect, and fixing it using Vetbond medical glue. The skin incision was also closed using Vetbond. A recovery cocktail consisting of sterile sodium chloride 0.9% saline with 1.25 mg.mL⁻¹ of atipamezole (Antisedan) and 0.01 mg.mL⁻¹ of buprenorphine (Vetergesic) was injected intraperitoneally into the mice (10 µL per g body weight), which were monitored and observed very carefully in an incubator at 28°C until recovered, after which they were returned to routine housing. All procedures involving animals were carried out by holders of Personal Licences (PIL) under a Project Licence (PPL, P8F273FDD, S.J.H.) shaped by ARRIVE and NC3Rs guidelines. The PPL was approved by the Animal Welfare and Ethical Review Body (AWERB) at King’s College London and granted by the Home Office under the Animals (Scientific) Procedures Act 1986 (ASPA) and Amendment Regulations 2012.

### μCT scanning and analysis

Eight weeks after transplantation, mouse head samples were collected. The hole head samples were fixed with 40 mL of 4% paraformaldehyde at room temperature overnight and washed three times with PBS for 15 min each the next day. The fixed samples were scanned using a μCT50 micro-CT scanner (Scanco). The specimens were immobilized in 19-mm scanning tubes using cotton gauze and scanned to produce voxel-size volumes of side length 10 μm using X-ray settings of 70 kVp and 114 μA and a 0.5-mm aluminium filter to attenuate harder X-rays. The scans were automatically scaled during reconstruction using the calibration object provided by the CT manufacturer, consisting of five rods of hydroxyapatite (HA) at densities of 0–790 mg HA cm–3, and the absorption values were expressed in Hounsfield Units (HU). The specimens were characterized using Parallax Microview software (Parallax Innovations), and 3D images were reconstructed from the raw μCT scanning data and analysed to quantify the area and volume of new bone regeneration. Background signals corresponding to soft non-mineralized tissue (<250 mg HA cm–3) and air were removed by thresholding, and mineralized tissue in the defect was considered to have a density of >500 mg HA cm^-3^. The 3D images were viewed and acquired using SCANCO Visualizer 1.1, and the virtual 2D sections were viewed using Fiji software.

### Cryosection preparation

Samples were decalcified overnight in 10% formic acid and then washed three times with PBS for 15 min each. Dehydration was carried out over two nights, first with 30% sucrose in water, and then with 30% sucrose and 30% optimal cutting temperature compound (OCT) in water. The samples were flash-frozen in cryomolds filled with OCT and stored at −80°C until they were sectioned. Mouse head samples were transversely sectioned at 12–14 μm in the anterior to posterior direction using a cryostat (Leica). Sectioned slices were stored at −20°C until stained.

### Histological staining

For Haematoxylin and Eosin (H&E) staining, the slides were prewarmed at room temperature for 10 min, then the OCT was dissolved with PBS for 5 min, twice. The slides were covered with Haematoxylin (Merck, MHS1) for 2 min, then rinsed twice with distilled water (10 s each), followed by ethanol 100% for 10 s. The slides were then covered with eosin (Merck, 318906) for 2 min, rinsed with distilled water (10 s), then dehydrated by a graded bath of ethanol and mounted using DPX (Dibutylphthalate Polystyrene Xylene, Merck, 06522) mounting medium. Movat’s pentachrome staining (Abcam, ab245884) was carried out according to the manufacturer’s instructions. The stained tissue sections were imaged with an RGB camera (Axiocam 208 color, Zeiss).

### Immunofluorescence staining of tissue sections

The slides were prewarmed at room temperature for 10 min, then the OCT was dissolved with PBS for 5 min, twice. The samples were permeabilized using 0.5% Triton X-100 in PBS for 5 min. The primary antibodies (human beta 2-microglobulin; Thermofisher, PA5142897, 1:100 (v/v); H3K14ac (Abcam, ab52946, 1:100 v/v); H3K9ac (Abcam, ab10812, 1:100 v/v); H3K27ac (Abcam, ab4729, 1:100 v/v) were diluted in a staining buffer (0.05% Triton X-100, 1% BSA in PBS) and left overnight at +4°C in a humid atmosphere. The samples were then washed with 0.1% Triton X-100 in PBS (3 × 5 min), then the secondary antibodies (Donkey anti-goat - AF488; Thermofisher, A-11055; 1:1000 v/v; Donkey anti-rabbit-AF555; Thermofisher, A-31572; 1:1000 v/v) were added to the sample in the staining buffer containing DAPI (1:1000 v/v) for 2 h at room temperature in the dark.After incubation with the secondary antibodies, the samples were washed with 0.1% Triton X-100 in PBS (3 × 5 min), and mounted with Fluoroshield (Merck, F6182). All types of staining were confirmed in tissues on at least three separate animals.

### Imaging and analysis of immunostained tissue sections

All images were acquired using an inverted spinning disk confocal microscope (Nikon, NIS-Elements Viewer 5.2 software), with a 60x magnification objective (in air) (NA=0.95; WD=200 µm). All the images were acquired with the following parameters: Z-step: 1 µm; DAPI laser power: 30%, exposure 500 ms; 488 laser power: 50%, exposure 700 ms; 555 laser power: 25%, exposure 500 ms. All imaging data were analysed using ImageJ software. The proportion of human cells (positive for human beta 2-microglobulin) was quantified by threshold analysis, as previously described^3,10^. Briefly, an area of stained control tissue (i.e., bone tissue or soft tissue) was selected and the highest intensity cellular signal of the marker of interest was determined. This intensity was used to threshold the sample tissue such that the remaining signal was of a higher intensity than that of the control tissue. The number of positive human cells could then be quantified based on the cellular signal. The nuclei of each human cell were then outlined and the intensity of H3K14ac quantified within the nuclei.

### Mechanical characterisation of PCL and PLGA films

Mechanical characterization was performed using a DMA Q8000 V7.5 Build 127 (TA instruments). Tests were run under at a fixed frequency of 1 Hz and an applied strain amplitude of 0.5%. All measurements were carried out within a controlled temperature range of 35–40 °C, ensuring thermal equilibration of each specimen before data acquisition.

### Statistical analysis

Statistical analysis was performed using GraphPad Prism 10. All data are plotted as mean, with error bars indicating standard deviation. Experimental data was analyzed comparing all conditions together using one-way ANOVA, with individual comparisons between conditions/regions of interest displayed on figures performed using uncorrected Welch’s two-tailed unpaired tests or uncorrected Fisher’s LSD test. Unpaired Welch’s t-tests were used to compare two-group datasets. Two-factors comparisons were analyzed with two-way ANOVA and uncorrected Fisher’s LSD test. Only the statistical values for the hypotheses being tested are shown; multiple comparison corrections were not applied. No randomization method was applied, nor blinding. The number of replicates is indicated in the figure or in the figure legends.

## Data availability statement

All data generated or analyzed during this study are included in this published article.

## Code Availability

Sequencing data is available using the following link and GEO number: GSE310737 (https://www.ncbi.nlm.nih.gov/geo/query/acc.cgi?acc=GSE310737)

## Author contributions

S.J.H. conceived the project and with P.B, P.T, S.J, B.M, X.C, D.F designed the experiments. P.B, P.T, S.J performed and analyzed experiments. Y.L, Y.T and D.F analyzed the genomic data. P.B, P.T and S.J.H wrote the manuscript. S.J, B.M, Y.L, Y.T, X.C, D.F and APK read, reviewed, edited and approved the manuscript. X.C, D. F and S.J. H supervised the project and S.J.H provided financial support for the project.

