## Supplementary materials for "Wnt-presenting materials sustain H3K14-acetylation in human skeletal stem cells for tissue engineering and bone repair"

### Supplementary Figures

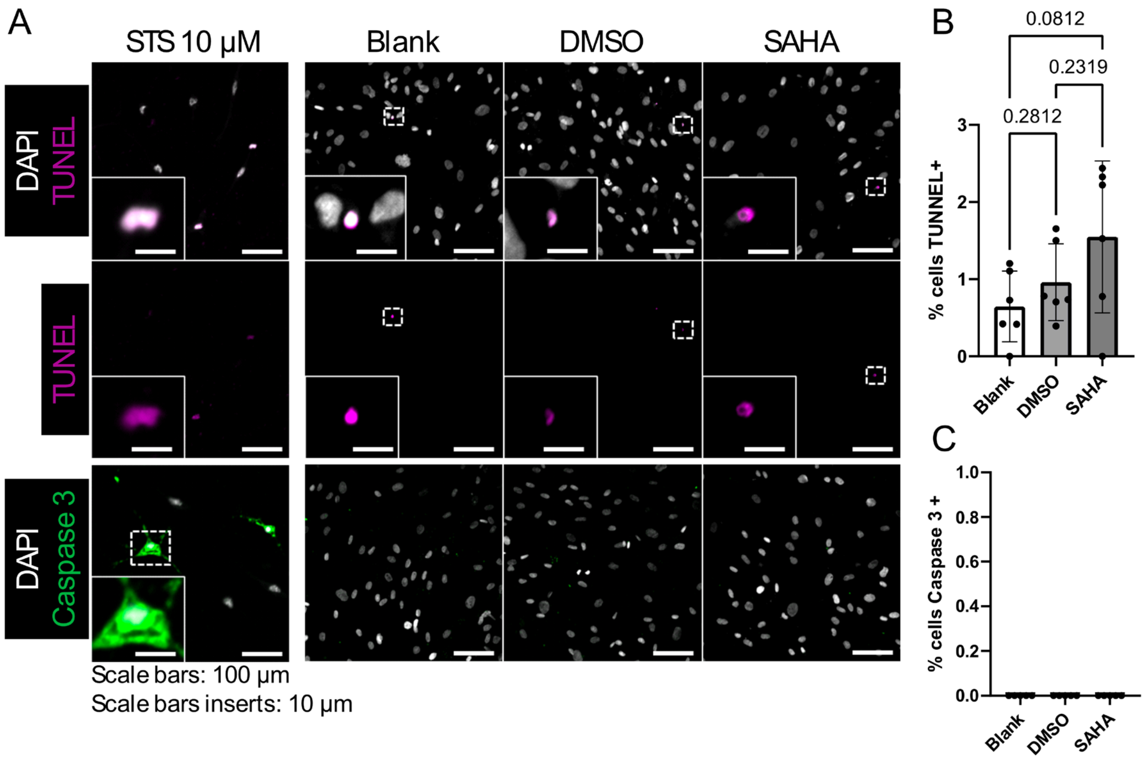


Supplementary figure 1 : SAHA treatment of hSSC in vitro does not induce cell death.
(A) Representative images and quantification of (B) TUNEL staining (purple) and (C) cleaved caspase 3 immunostaining (green) in the WIOTM after 7 days of culture under three conditions: osteogenic media alone (Blank), osteogenic media + DMSO, and osteogenic media + SAHA (1 µM). Staurosporine 10 µM was added 6h before the end of the experiment and used as a positive control of cell death. The data represents the mean ± s.d. Statistical analysis: ANOVA followed by unpaired t test with Welch's correction. N=6 WIOTM, n ≥ 97 cells per WIOTM.

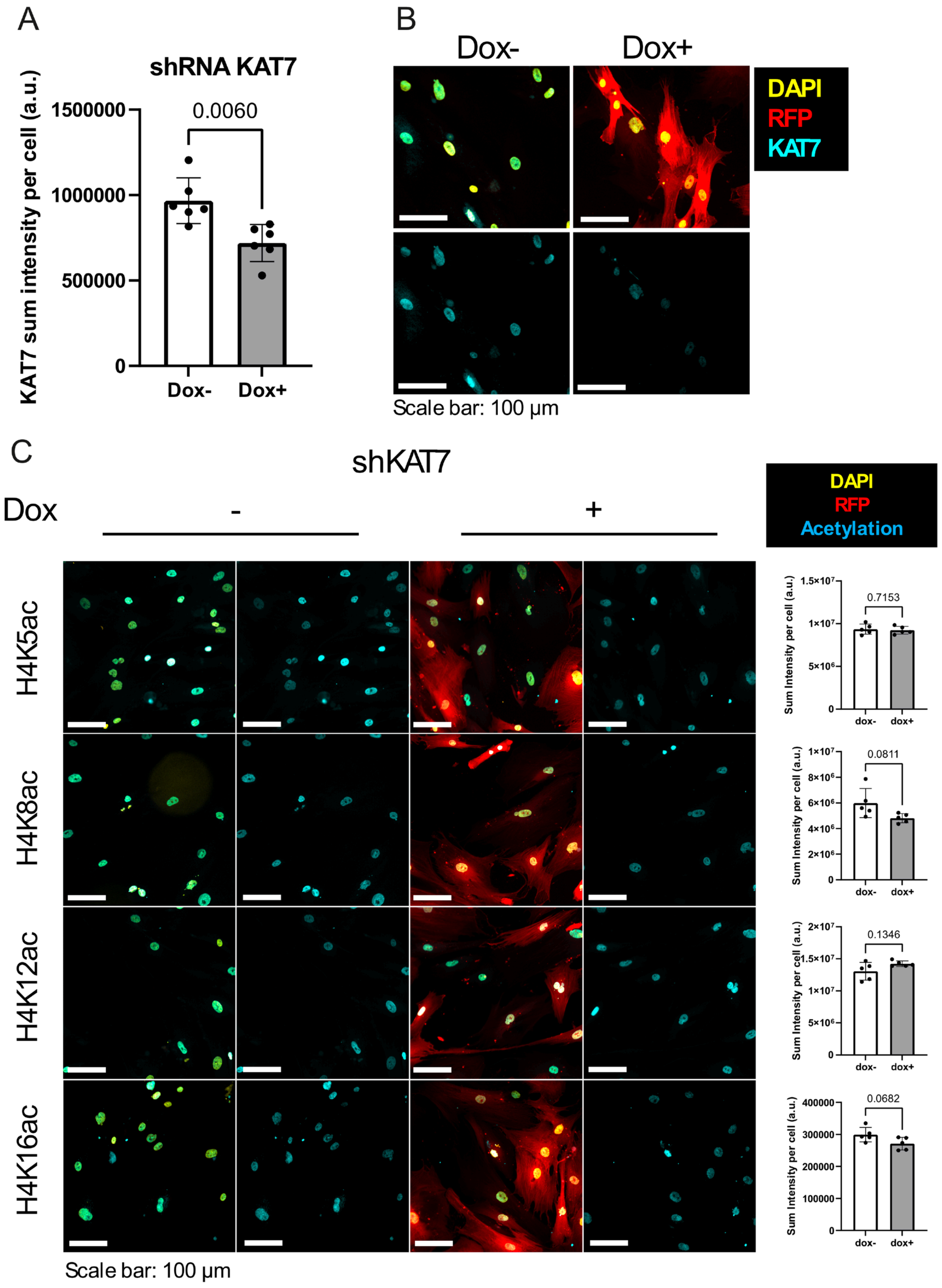

A
shRNA KAT7
KAT7 sum intensity per cell (a.u.)
1500000
1000000
500000
0
0.0060
Dox-
Dox+
B
Dox-
Dox+
DAPI
RFP
KAT7
Scale bar: 100 µm
C
shKAT7
Dox
-
+
DAPI
RFP
Acetylation
H4K5ac
H4K8ac
H4K12ac
H4K16ac
Sum Intensity per cell (a.u.)
0.7153
0.0811
0.1346
0.0682
dox-
dox+
Scale bar: 100 µm

Supplementary figure 2: Kat7 knock-out does not affect H4 acetylation.

(A) Quantification and (B) representative images of Kat7 (cyan) immunostaining in hSSCs after transduction of a doxycycline inducible shRNA targeting Kat7 and 7 days of induction. Induction of the shRNA with doxycycline treatment also induces the expression of the red fluorescent protein (RFP) shown in red. The data represents the mean ± s.d. Statistical analysis: Uncorrected Fisher's LSD tests. N=6 biological replicates, n ≥ 21 cells. (C) Representative images and quantification of the expression of a panel of H4 acetylation in hSSCs transduced with shRNA targeting *Kat7* after 7 days of induction. Acetylation marks are shown in blue, RFP in red. The data represents the mean ± s.d. Each data point represents a biological replicate. Statistical analysis: Uncorrected Fisher's LSD tests. N=5 biological replicates, n ≥ 19 cells.

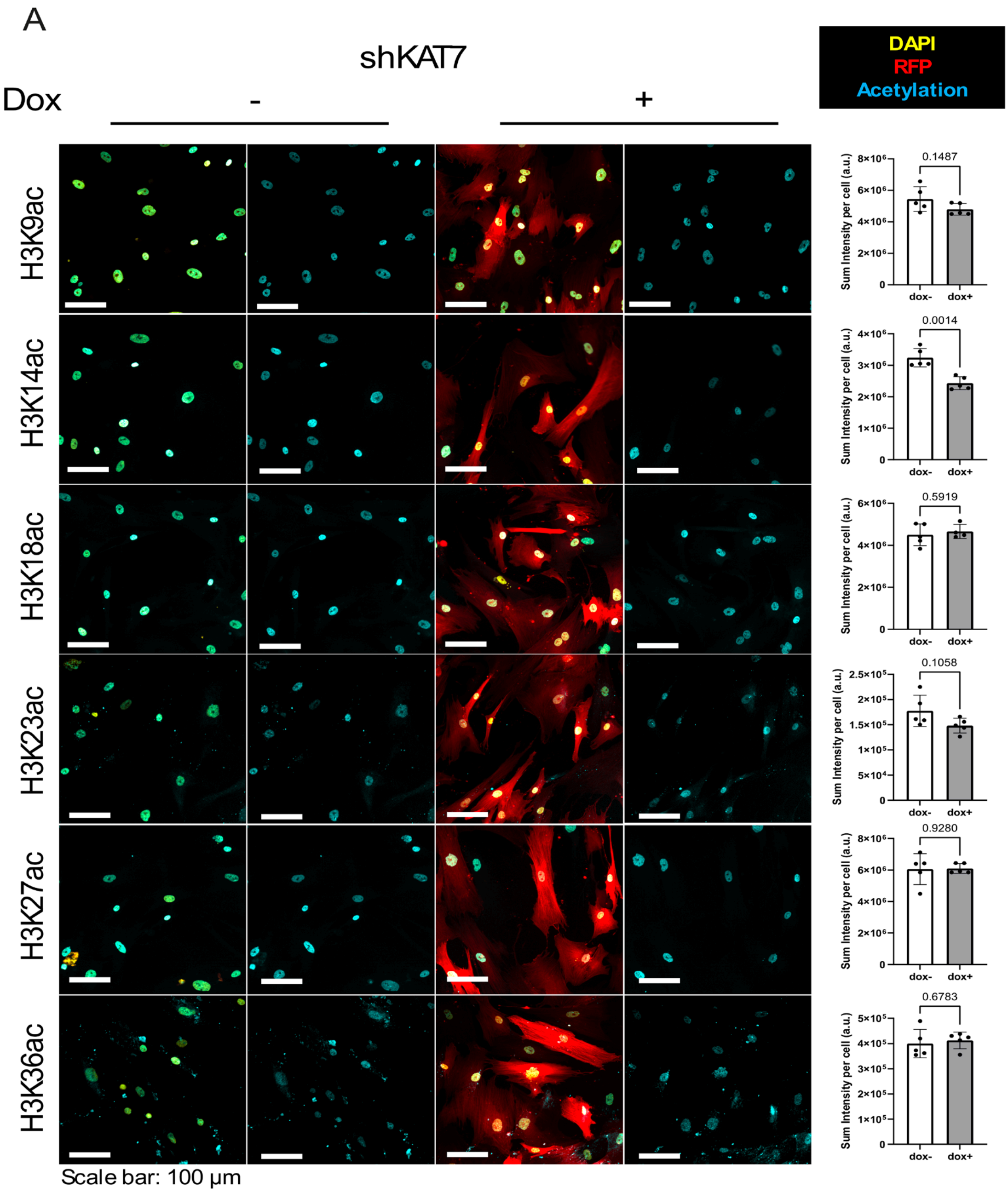


Supplementary figure 3: Kat7 knock-out only affects H3K14ac.
(A) Representative images and quantification of the expression of a panel of H3 acetylation in hSSCs transduced with shRNA targeting *Kat7* after 7 days of induction. Acetylation marks are shown in blue, RFP in red. The data represents the mean ± s.d. Each data point represents a biological replicate. Statistical analysis: Uncorrected Fisher's LSD tests. N=5 biological replicates, n ≥ 23 cells.

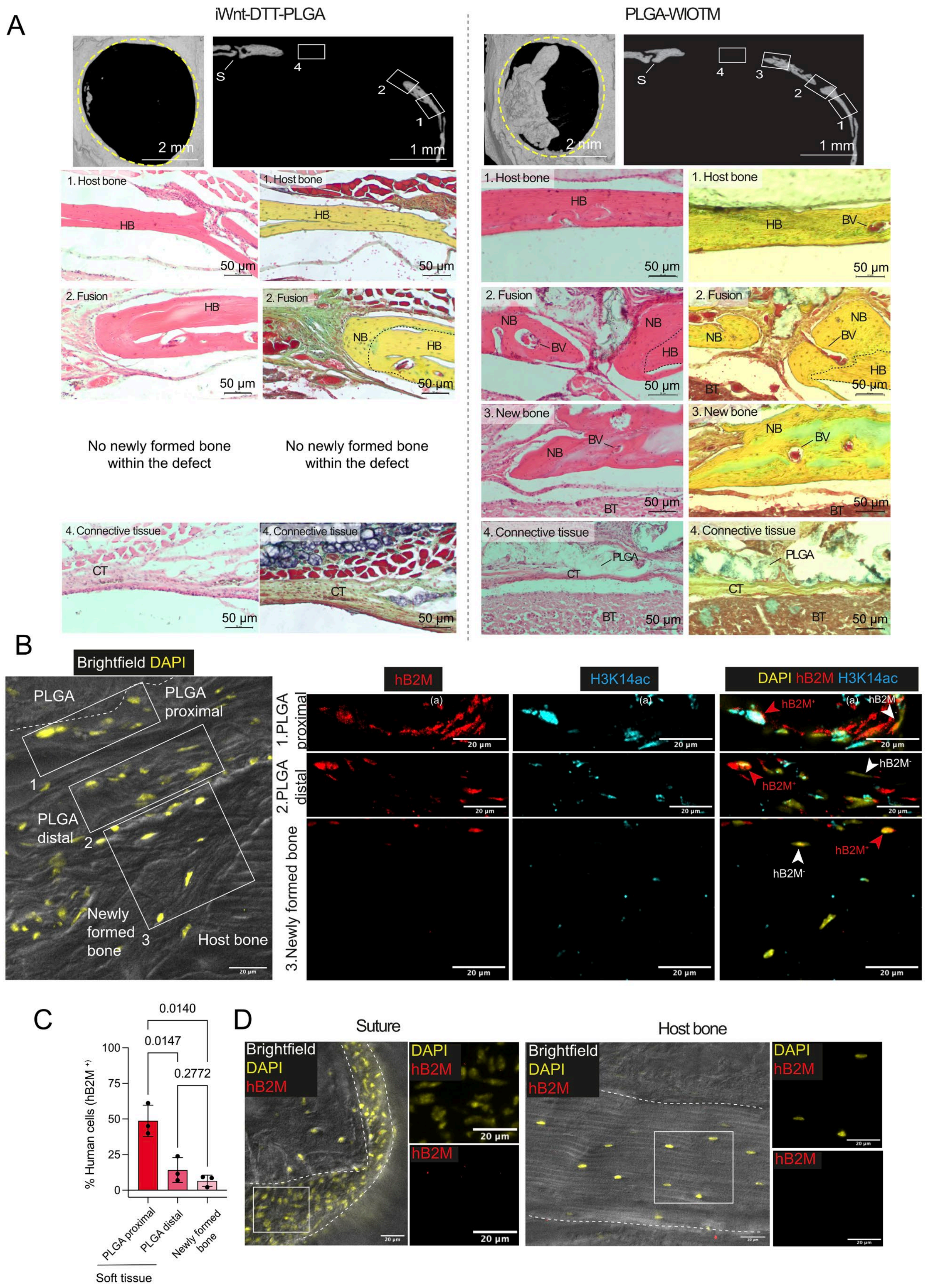
A
iWnt-DTT-PLGA
PLGA-WIOTM
S
1
2
3
4
2 mm
1 mm
1. Host bone
2. Fusion
3. New bone
4. Connective tissue
HB
NB
BV
BT
CT
PLGA
50 µm
No newly formed bone within the defect
B
Brightfield DAPI
PLGA
PLGA proximal
PLGA distal
Newly formed bone
Host bone
hB2M
H3K14ac
DAPI hB2M H3K14ac
1.PLGA proximal
2.PLGA distal
3.Newly formed bone
hB2M+
hB2M-
20 µm
C
% Human cells (hB2M+)
0.0140
0.0147
0.2772
PLGA proximal
PLGA distal
Newly formed bone
Soft tissue
D
Suture
Host bone
Brightfield
DAPI
hB2M

Supplementary figure 4: Histological characterization of the bone defects.
(A) Representative µCT images with top view and frontal digital sections of defects covered by iWnt-DTT-PLGA or PLGA-WIOTM. The dashed yellow circle indicates the edges of the drilled defects. S: sagittal suture of the calvaria. The numbered squares (1-4) indicate the enlarged area where representative images of H&E and Movat's pentachrome staining have been taken. HB: host bone, BV: blood vessel, NB: newly formed bone, HB: host bone, BT: brain tissue, CT: connective tissue. The dashed black line indicates the limit of the host bone (i.e., the edge of the drilled defect). (B) Representative images of the newly formed bone overlaid by the soft tissue, and the PLGA bandage. The dashed line indicates the limit of the PLGA. The numbered (1-3) white squares indicate the enlarged area on the right panel. The human marker (human beta-2 microglobulin (hB2M)) is displayed in red and indicates cells of human origin, based on tissue-specific threshold. The red and white arrowheads indicate positive ($hB2M^+$) and negative ($hB2M^-$) cells, respectively. H3K14ac is displayed in blue with the same threshold in all images. (C) Proportion of human cells ($hB2M^+$) in each tissue class. Data are presented as mean ± S.D. Statistical analyses: uncorrected ANOVA test. N=3 animals, n = 194 cells. (D) Representative images of immunostaining of host tissue (suture and host bone) where no human cells (labelled by hB2M) were found. The dashed lne indicates the limit of the sagittal suture (left panel), or the host bone (right panel).

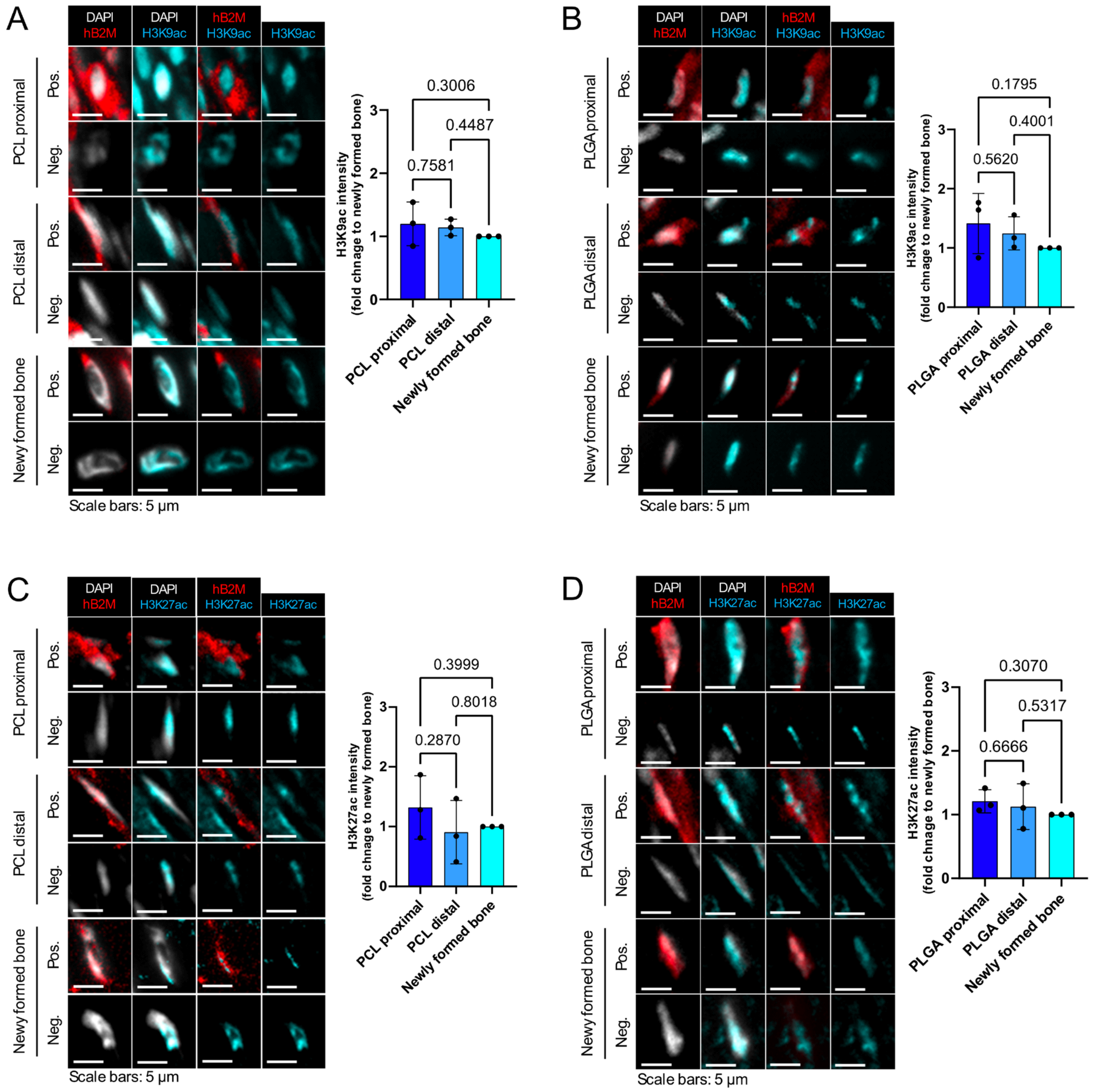


Supplementary figure 5: Localized Wnt-3a signaling on biodegradable polymers does not polarize H3K9ac and H3K27ac in transplanted hSSCs in a critical size defect model in mice calvaria.

(A) Representative images and quantification of tissue sections immunostained with H3K9ac and human beta-2 microglobulin. PCL proximal (i.e., close to the PCL), PCL distal (i.e., at distance from the PCL, close to the newly formed bone); newly formed bone (i.e., within the newly formed bone tissue). The human marker (human beta-2 microglobulin) is displayed in red and indicates cells of human origin, based on tissue-specific threshold (native bone). H3K9ac is displayed in blue with the same threshold in all images. H3K9ac intensity is quantified in cells of human origin in each tissue class. N = 3 animals, n = 149 cells. (B) Similar to (A) but here the calvarial defect was covered with a PLGA WIOTM bandage. N = 3 animals, n = 159 cells. (C) Similar to (A) but sections were immunostained with with H3K27ac (blue) and human beta-2 microglobulin (Red). N = 3 animals, n = 130 cells. (D) Similar to (C) but here the calvarial defect was covered with a PLGA WIOTM bandage. N = 3 animals, n = 137 cells. All data are presented as mean ± S.D. Statistical analyses: ANOVA followed by uncorrected Fisher's LSD.

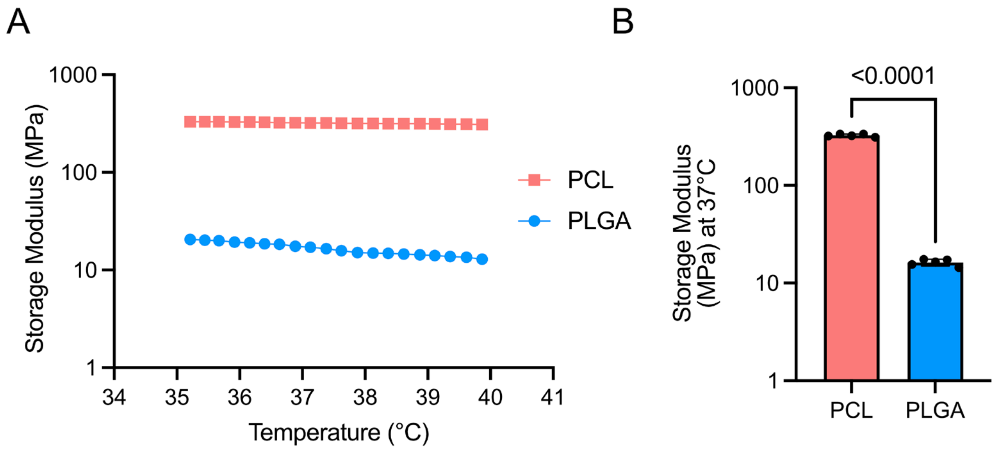


Supplementary figure 6: Mechanical properties of PCL and PLGA films

(A) Temperature ramp and measure of the storage moduli of polycaprolactone (PCL) and poly(lactic-co-glycolic acid) (PLGA) films (1 Hz and 0.5% strain) within a temperature range of 35-40 °C.(B) Storage moduli at 37 °C for PCL and PLGA films. Data are presented as mean ± S.D. N=5 films per group. Statistical analysis: unpaired Welch's t-test.

| Fig3C-D | DAPI | H3K14ac | H3K9ac | RNApolS2P | OPN:CDH13 |
|---|---|---|---|---|---|
| DMSO | <0.0001 | <0.0001 | 0.0022 | <0.0001 | 0.0061 |
| SAHA | 0.0083 | 0.3802 | 0.9194 | 0.008 | 0.4987 |

Supplementary table 1: p-values after one sample t-test from figure 3C and D. Camparison to the theorical value of 1

| Fig4B & D | H3K14ac | | OPN:CDH13 | |
|---|---|---|---|---|
| | shScramble | shKAT7 | shScramble | shKAT7 |
| dox- | <0.0001 | 0.0019 | 0.0032 | 0.0143 |
| dox+ | 0.0257 | 0.9712 | 0.0017 | 0.0206 |

Supplementary table 2: p-values after one sample t-test from figure 4B and D. Camparison to the theorical value of 1
